# Single-nucleus profiling reveals age-associated remodeling opposed by parity in the postmenopausal human ovary

**DOI:** 10.64898/2026.05.11.724381

**Authors:** Josef Byrne, Nicolas Martin, Mark A. Watson, Bikem Soygur, Pooja Raj Devrukhkar, Tommy Tran, Elisheva Shanes, Mary Ellen G. Pavone, Francesca E. Duncan, Birgit Schilling

## Abstract

The postmenopausal ovary is commonly viewed as a largely quiescent organ, yet its cellular biology remains incompletely characterized. We generated a single-nucleus atlas of the aging postmenopausal human ovary comprising 439,011 nuclei across 64 ovarian samples from 28 donors. We resolved 37 cell states, revealing extensive stromal, vascular, and immune heterogeneity. We developed an interactive data explorer to facilitate interrogation of cell annotations and gene expression across the atlas. Aging was associated with expansion of stress-associated stromal states, vascular and immune depletion, and a shift toward steroidogenic and androgen-responsive programs. Several major age-associated compositional shifts were supported in an external GTEx ovary bulk RNA-seq cohort, validating relevance. Parity (1–3 versus 0 live births) was associated with transcriptional and compositional patterns broadly opposing age-associated remodeling. Our findings establish the postmenopausal ovary as an actively remodeled aging tissue and link obstetric history to molecular and cellular signatures detectable later in life.

## Main

The ovary is one of the first major organ systems in women to show functional decline, with reproductive potential declining well before menopause^1,2^. The consequences of ovarian aging extend beyond fertility, contributing to endocrine dysfunction and adverse cardiometabolic, skeletal, and overall health outcomes^2–5^. Yet, ovarian aging is still often framed primarily through follicle depletion and endocrine failure rather than as a broader process involving age-associated remodeling of the ovarian tissue microenvironment. Because women in many countries now live decades beyond menopause, understanding the molecular and cellular composition of the postmenopausal ovary is essential for determining how this organ continues to influence local signaling, systemic hormone exposure, and gynecologic disease risk after the final menstrual period.

Recent cellular atlases have begun to refine the view of the aging ovary as a structurally and molecularly heterogeneous tissue. Single-cell and single-nucleus studies indicate that human ovarian aging involves coordinated changes across stromal, vascular, immune, and follicular compartments, and that these changes are spatially organized rather than uniform^6–12^. In postmenopausal ovarian tissue, stromal cells dominate whereas follicular elements are profoundly depleted, suggesting that later ovarian aging may be shaped less by oocyte-centered processes and more by stromal states, microvascular support, and intercellular signaling. Consistent with this shift, a growing body of work in murine models, non-human primates, and humans has implicated the stromal microenvironment as a major locus of ovarian aging, with reproductively aged ovaries accumulating collagen-rich fibrotic foci, becoming mechanically stiffer, undergoing broad extracellular-matrix remodeling, and acquiring phenotypes associated with cellular senescence across histologic, biomechanical, and proteomic studies^13–21^. The immune and vascular compartments also appear relevant, although their contributions remain less defined. Human studies highlight macrophage-state remodeling during reproductive aging together with changes in vascular cell abundance in comparisons of young and reproductively aged ovaries^6,11^, whereas mouse studies emphasize broader immune accumulation, inflammatory niche formation, and vascular decline as a potential driver of ovarian aging^22–28^. Together, these observations suggest that a central unresolved question is which multicellular biological programs persist in the residual ovary after reproductive cessation^29^.

Another unresolved question is whether the postmenopausal ovary remains steroidogenically active and biologically influential after reproduction ceases. Studies measuring ovarian-vein hormone levels and examining women following oophorectomy support a continued ovarian contribution to androgen production after menopause^30,31^. Furthermore, recent mass spectrometry-based analyses suggest that circulating testosterone in older women may increase during later menopausal stages^32,33^.

Beyond chronological aging, reproductive history may also leave durable biological signatures in the postmenopausal ovary. Epidemiologic studies suggest that parity (lifetime live birth count) is linked to menopause timing, biological aging trajectories, and late-life survival in nonlinear ways, with moderate parity often associated with more favorable outcomes than nulliparity or high parity^34–37^. Experimental and mechanistic studies further suggest that pregnancy can induce durable immune and epigenetic changes, and that microchimeric cells may persist for decades after gestation^38,39^. Whether these long-lived effects are detectable as cellular or molecular signatures in the postmenopausal ovary remains unknown.

Here, we generated a single-nucleus RNA-seq atlas of the postmenopausal human ovary and integrated compositional, transcriptional, and cell–cell communication analyses to characterize ovarian remodeling associated with aging and parity. We identified coordinated stromal–immune remodeling, vascular attrition with residual microvascular remodeling, and expansion of steroidogenic stromal programs accompanied by androgen-responsive transcriptional changes. We further observed broad opposition between age- and parity-associated molecular and compositional signatures. Together, these findings establish the postmenopausal ovary as an active aging tissue and an important system for studying how obstetric history intersects with core aging-associated programs, while providing a resource to inform future studies of ovarian aging, women’s health, and geroscience.

## Results

### Single-nucleus profiling defines a cellular atlas of the postmenopausal human ovary

To define the cellular landscape of the postmenopausal human ovary and examine its remodeling with age, we performed droplet-based single-nucleus RNA-sequencing (snucRNA-seq) on 64 frozen ovarian tissue pieces containing cortical and medullary regions from 28 donors undergoing oophorectomy (**Fig. 1a**). Donors ranged from 50 to 84 years of age (**Fig. 1b**) and had parity values of 0–5 (**Fig. 1c**). The majority of donors had their ovaries removed for benign gynecologic surgical indications (**Fig. 1d**). In this cohort, consistent with reported population-level associations, parity was positively associated with urinary incontinence and pelvic organ prolapse and inversely associated with uterine malignancy (**Extended Data Fig. 1**)^40,41^. All ovarian tissue used for downstream analysis was confirmed by a gynecologic pathologist to be free of premalignant and malignant ovarian pathology.

**Fig 1.**
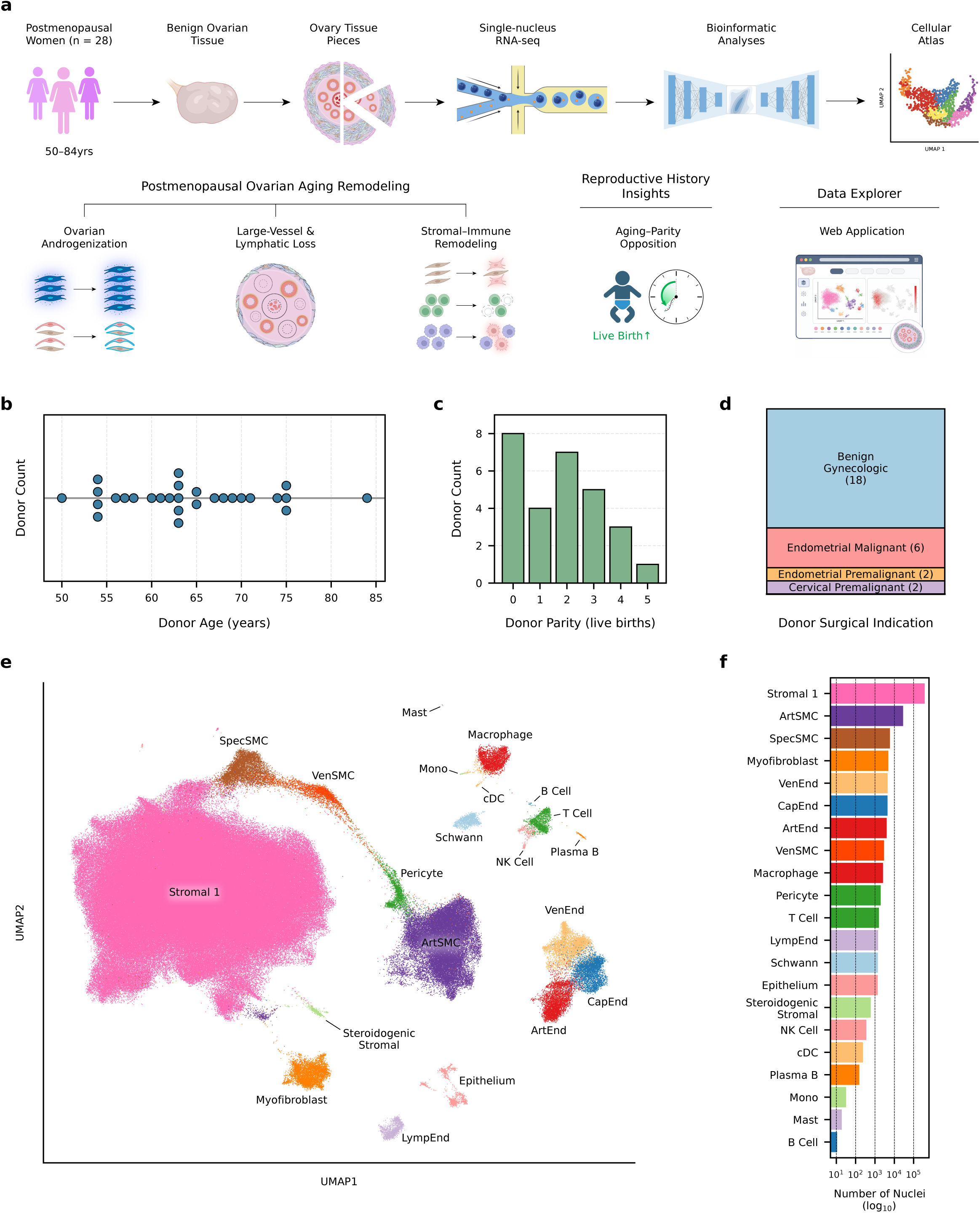
Single-nucleus RNA-seq atlas of the aging human postmenopausal ovary. a, Schematic overview of study design, experimental workflow, and primary study findings. Illustration was created using BioRender.com. **b,** Age distribution of the cohort. Each dot represents one donor. **c,** Parity (lifetime number of live births) distribution of the cohort. **d,** Surgical indications for ovary procurement. **e,** Uniform manifold approximation projection (UMAP) of annotated nuclei in the atlas. **f,** Number of nuclei per annotated cell type shown on a log_10_ scale. ArtSMC, arterial smooth muscle; VenSMC, venous smooth muscle; SpecSMC, specialized smooth muscle; ArtEnd, arterial endothelial; VenEnd, venous endothelial; CapEnd, capillary endothelial; LympEnd, lymphatic endothelial; Mono, monocyte; cDC, conventional dendritic cell.

Following preprocessing, quality control, and single-cell variational inference (scVI)-based integration, we recovered 439,011 high-quality nuclei that resolved into 21 coarse cell types (**Fig. 1e**; see Methods). Canonical marker expression supported these annotations, and the integrated embedding showed no overt gross segregation by donor, processing batch, or non-ovarian malignancy status (**Extended Data Fig. 2**). The atlas was dominated by a large stromal cell population (∼85% of nuclei; annotated as Stromal 1), but also captured diverse endothelial, smooth muscle/perivascular, immune, epithelial, Schwann, myofibroblast, and steroidogenic stromal populations (**Fig. 1f**). This stromal predominance is consistent with the follicle-depleted architecture of the postmenopausal ovary and illuminates the residual stroma as a major cellular component of later-life ovarian remodeling.

### Transcriptional characterization of ovarian cell type compartments

To further resolve transcriptional heterogeneity within the atlas, we reanalyzed each major compartment independently using the same dimensional reduction, clustering, and annotation framework applied at the atlas level, identifying 37 fine cell states across stromal, endothelial, mural, epithelial, Schwann, and immune compartments (**Fig. 2a**; **Extended Data Fig. 3**; Supplementary Table 2). This higher resolution analysis recovered expected lineage-resolved states while revealing rich stromal heterogeneity. Notably, no distinct granulosa or oocyte populations were recovered, consistent with the marked depletion of follicles after menopause. This cellular landscape highlights stromal, vascular, immune, epithelial, and neural-support compartments as the predominant cellular niches of the postmenopausal ovary.

**Fig 2.**
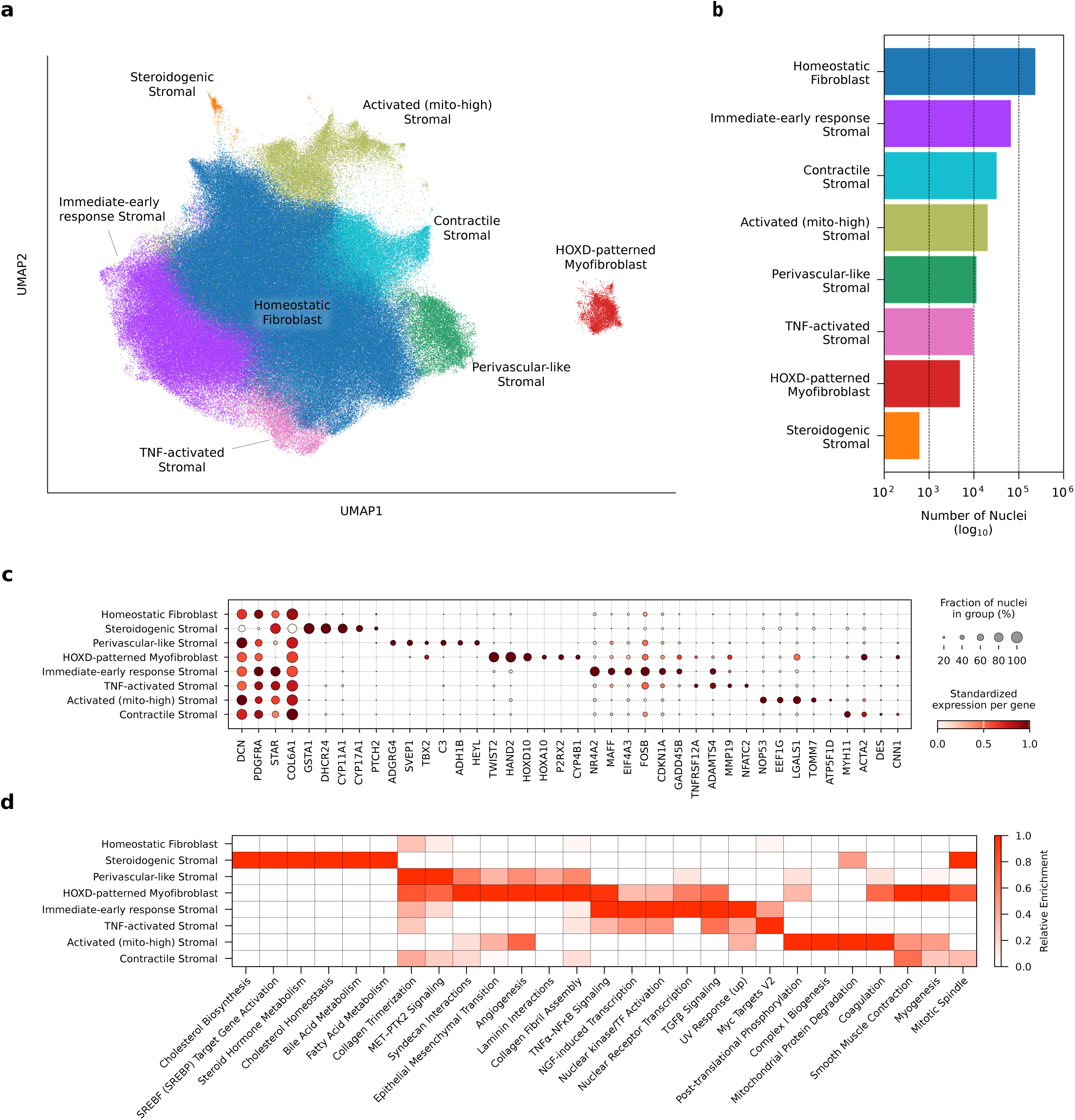
Ovarian stromal compartment characterization. a, UMAP showing annotated stromal states observed in the postmenopausal ovary. **b,** Horizontal bar plot showing number of nuclei per annotated stromal state on a log_10_ scale. **c,** Representative marker genes distinguishing each stromal state. Gene expression was scaled across stromal states, and dot size indicates the fraction of nuclei in each stromal state with detectable expression. **d,** Descriptive pathway heatmap showing curated MSigDB Hallmark and Reactome pathways distinguishing stromal states. Heatmap cells are colored by relative enrichment after per-pathway min–max scaling.

Within the stromal compartment, Stromal 1, myofibroblast, and steroidogenic stromal populations separated into eight transcriptionally distinct states spanning homeostatic, steroidogenic, contractile, vascular-adjacent, and stress-associated programs (**Fig. 2**). A large homeostatic fibroblast population formed the core of the stromal compartment, expressing *DCN*, *PDGFRA*, *STAR*, and *COL6A1* without a sharply restricted state-defining marker program, consistent with a baseline fibroblast identity. In contrast, a discrete steroidogenic stromal state was distinguished by *GSTA1*, *DHCR24*, *CYP11A1*, and *CYP17A1* and enriched for cholesterol biosynthesis and steroid hormone metabolism pathways. This profile is suggestive of a non-follicular theca interna-like program. We also identified a contractile stromal state marked by *MYH11*, *ACTA2*, *DES*, and *CNN1*, together with smooth muscle contraction and myogenic pathway enrichment, consistent with a contractile stromal program with theca externa-like features.

A perivascular-like stromal population was distinguished by *ADGRG4*, *SVEP1*, *C3*, *ADH1B*, and *HEYL*, consistent with a vascular-adjacent stromal program (**Fig. 2**). We also identified a transcriptionally distinct HOXD-patterned myofibroblast population marked by *TWIST2*, *HAND2*, *HOXD10*, *HOXA10*, *P2RX2*, and *CYP4B1*. This state remained distinct from the canonical contractile stromal state and was enriched for epithelial–mesenchymal transition, angiogenesis, and collagen-associated extracellular-matrix remodeling pathways, including collagen fibril assembly. This population was also the state with the highest score for the recently defined BuckSenOvary senescence-associated signature^16^ (**Extended Data Fig. 4a**; see Methods), a 32-gene program derived from p16-associated postmenopausal ovarian regions showing inflammatory and extracellular-matrix remodeling and increased collagen architectural complexity. Together, these observations associate the HOXD-patterned myofibroblast population with fibro-inflammatory senescence-associated stromal microenvironments within the postmenopausal ovary.

The stromal compartment also contained multiple transcriptionally distinct stress-associated states rather than a single generic activated population. The most prominent of these was an immediate-early response (IER) stromal state marked by *FOSB*, *NR4A2*, *CDKN1A*, and *GADD45B* and enriched for TNFα–NFκB, TGFβ, and UV response pathways, consistent with engagement of immediate-early and stress response signaling (**Fig. 2**). In parallel, a rarer TNF-activated stromal state was distinguished by *TNFRSF12A*, *MMP19*, and *NFATC2*, whereas an activated mito-high stromal state was enriched for mitochondrial and translational genes, including *TOMM7*, *ATP5F1D*, and *EEF1G*, together with mitochondrial and proteostatic pathways. Although the mito-high state passed mitochondrial quality control thresholds, its expression profile suggests that it may reflect either an activated *in situ* stromal state or partial sensitivity to tissue handling.

Beyond the stromal compartment, finer annotation resolved heterogeneity within vascular, immune, epithelial, and neural-support populations. Endothelial states exhibited distinct arterial, capillary, venous, and lymphatic identities, while mural populations separated into pericytes and transcriptionally distinct smooth muscle states, including arterial and venous smooth muscle populations (**Extended Data Fig. 3a–d**). Immune subclustering resolved both innate and adaptive populations, including distinct antigen-presenting and lymphocyte subsets (**Extended Data Fig. 3e,h**). Within the epithelial compartment, mesothelial cells were distinguished from Müllerian epithelial states with differing *PAX2* expression and a separate stress-response secretory program (**Extended Data Fig. 3f,i**). Schwann cells further separated into states expressing myelinating or non-myelinating programs, extending the atlas to glial populations associated with ovarian innervation (**Extended Data Fig. 3g,j**).

Additional specialized states were also identified beyond the canonical cell types, including a small endothelial population defined by *SERPINE2*, *TIMP2*, *RNASE1*, and *CAVIN1*, suggestive of a specialized vascular-wall endothelial program associated with endothelial homeostasis, remodeling control, and caveolar signaling^42–45^ (**Extended Data Fig. 4b,c**). We also identified a distinct GPNMB+ macrophage population marked by *GPNMB*, *APOE*, *IL4I1*, and the fusion-associated gene *DCSTAMP*, consistent with features of ovarian multinucleated giant cells that have been reported to increase with reproductive aging in mice and non-human primates^13,26,46^ (**Extended Data Fig. 4d,e**). Together, these findings show that the postmenopausal ovary retains substantial multicellular heterogeneity despite profound follicular depletion.

To facilitate additional exploration and reuse of the atlas, we provide an interactive data explorer (https://huggingface.co/spaces/josef-byrne/postmenopausal-ovary-atlas-browser) for examining cell annotations and gene expression patterns across ovarian cell states (Supplementary Fig. 12; see Code availability).

### Cell type compositional analysis of ovarian aging

To assess how ovarian cell type composition varies with age, we modeled relative cell type abundance across the atlas using a Bayesian compositional framework that included biological and technical covariates, including a sample-level StressScore (see Methods). This analysis identified four coherent features of postmenopausal ovarian aging: expansion of steroidogenic and contractile stromal populations, reduction of large-vessel and lymphatic-associated cell types, contraction of several immune populations, and a shift from homeostatic toward stress-associated stromal states (**Fig. 3a,b** - left columns; Supplementary Fig. 5; Supplementary Table 4). In our snucRNA-seq atlas, homeostatic fibroblasts declined with age whereas immediate-early response (IER) and TNF-activated stromal states increased, consistent with a shift toward stress-associated stromal programs. Separately, steroidogenic stromal and contractile stromal populations also expanded with age. Outside the stromal compartment, venous and lymphatic endothelial cells, vascular smooth muscle populations, and several immune populations, notably CD4+ T cells, plasma B cells, and cDC2, were depleted. This coordinated pattern suggests that postmenopausal aging involves remodeling of the stromal microenvironment together with its vascular and immune compartments, rather than changes confined to a single cellular lineage.

**Fig 3.**
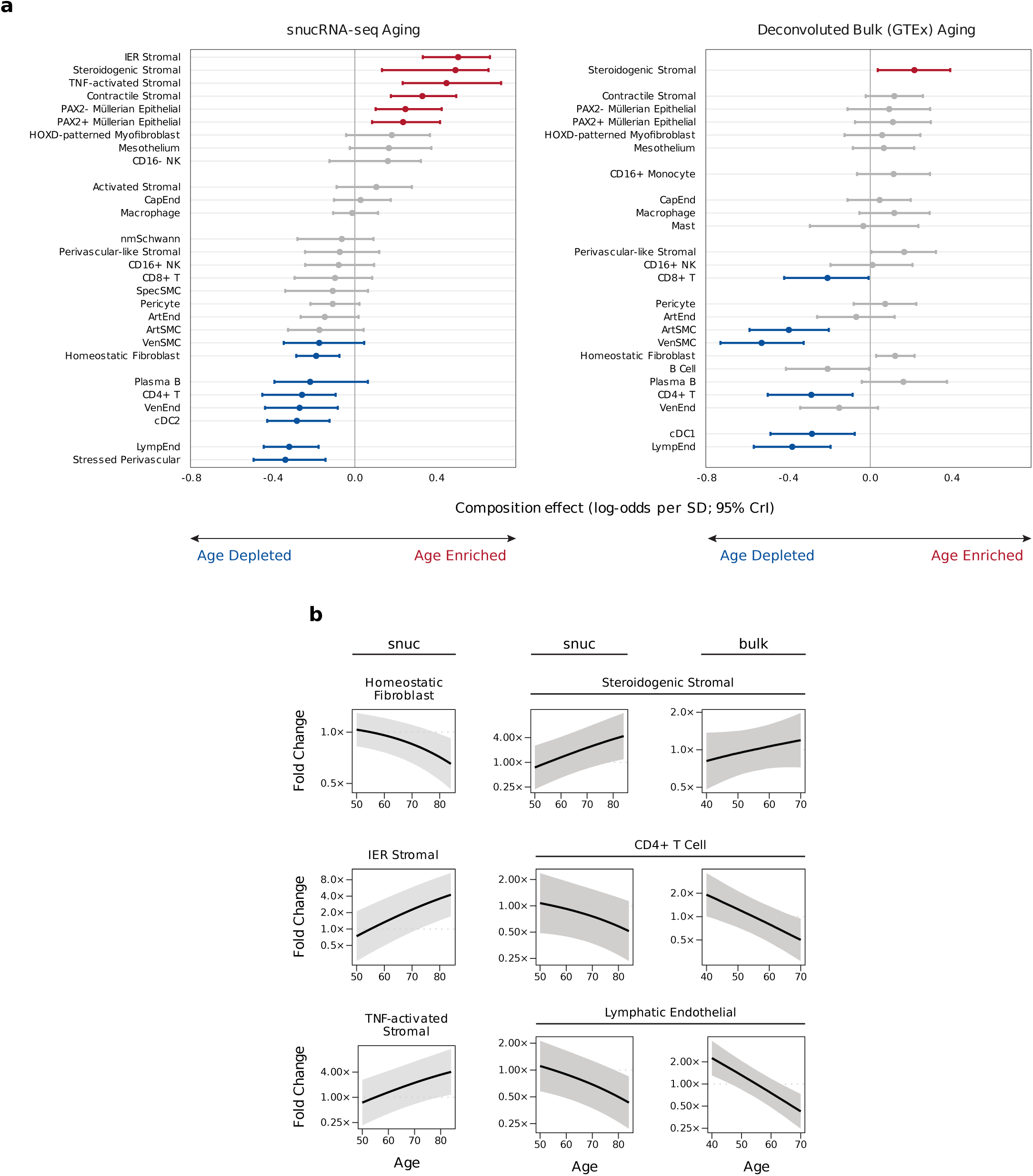
Cell type compositional analysis of snucRNA-seq and deconvolved bulk. a, Forest plots showing age-associated changes in cell type abundance in the snucRNA-seq atlas and deconvolved bulk GTEx RNA-seq data. Cell types not represented in one of the two analyses were omitted from that panel for clarity (see Methods). Points show posterior median composition effects on the log-odds (logit) scale per one-standard-deviation increase in age within each cohort. Error bars indicate 95% credible intervals. Colored entries indicate significant effects (Bayesian FDR ≤ 0.05) based on posterior evidence for an absolute composition effect exceeding 0.1 logit units. **b,** Model-predicted fold changes in proportion with age for representative cell types in snucRNA-seq and deconvolved bulk data, shown relative to age 55 years. Lines represent posterior median fold changes and ribbons represent propagated 95% credible intervals in fold-change space. IER, immediate-early response.

To test whether these age-associated compositional shifts generalized beyond the snucRNA-seq cohort, we deconvolved GTEx ovary bulk RNA-seq from donors enriched for likely postmenopausal status (n=104)^47^ using the atlas generated in this study as a reference and repeated cell type compositional modeling (see Methods). Across GTEx donors, deconvolved compositions spanned a broad continuum from homeostatic fibroblast-rich to more vascular/perivascular-rich profiles, consistent with sampling variation along a putative ovarian axis from stromal-dense, cortex-enriched tissue to vascularized, medulla-enriched tissue (**Extended Data Fig. 5a**). Deconvolution quality-control metrics supported interpretability of the deconvolved cell type profiles (**Extended Data Fig. 5b**; Supplementary Fig. 3; Supplementary Table 3; see Methods). Compositional modeling of the deconvolved GTEx cohort recapitulated several key age-associated trends, including enrichment of steroidogenic stromal cells and depletion of endothelial, smooth muscle, and T cell populations (**Fig. 3a,b**; Supplementary Fig. 6). Stress-associated stromal states were excluded from deconvolution because they did not yield sufficiently distinct bulk profiles, likely contributing instead to the broader homeostatic fibroblast fraction, which directionally increased with age in GTEx despite declining in the snucRNA-seq atlas. These findings provide external support for several major age-associated compositional trends detected in the atlas, while also illustrating the influence of tissue sampling and resolution on inferred ovarian cellular composition.

Although the donors of our cohorts were predominantly White, Black donors were represented in the snucRNA-seq atlas and GTEx bulk data (snucRNA-seq: 4 of 28 donors; GTEx: 14 of 104 donors). Two stromal cell types showed notable concordance in race-associated compositional differences across cohorts. In both datasets, Black donors showed higher relative abundance of steroidogenic stromal and contractile stromal populations, whereas most other shifts were not concordant across cohorts. However, given the limited number of Black donors, these race-associated differences should be considered exploratory (**Extended Data Fig. 6a,b**).

Across donors more broadly, steroidogenic stromal and contractile stromal abundances were strongly correlated across all donors in both snucRNA-seq and deconvolved GTEx data, consistent with linked abundance changes in these two stromal populations (**Extended Data Fig. 6c**). Together with the transcriptional resemblance of steroidogenic stromal and contractile stromal populations to theca interna-like and theca externa-like programs, this raises the possibility that these populations represent paired theca-like stromal states persisting despite the absence of detectable follicular granulosa and oocyte populations in the postmenopausal ovary. The presence of these theca-like states may provide a cellular basis for continued ovarian steroidogenic activity despite profound follicular depletion.

### Neighborhood-level analysis of immune cell abundance

To determine whether age-associated immune compositional changes extended to heterogeneity within annotated cell types, we applied Milo^48^, a graph-based differential abundance framework that tests partially overlapping transcriptional neighborhoods. We focused on the immune compartment because it exhibited cell type compositional differences and contained sufficient nuclei for neighborhood-level analysis (see Methods). After applying neighborhood-level quality control filters to exclude donor-dominated neighborhoods, we identified age-associated depletion of neighborhoods belonging to macrophage, CD8+ T, and CD4+ T cell populations (**Fig. 4a–c**; Supplementary Table 5). While the T cell changes were directionally concordant with the cell type compositional analysis, macrophage neighborhoods showed heterogeneous age-associated changes across the macrophage cluster, revealing intra-cell-type compositional structure not captured by the annotated cell type abundances (**Fig. 4c**).

**Fig 4.**
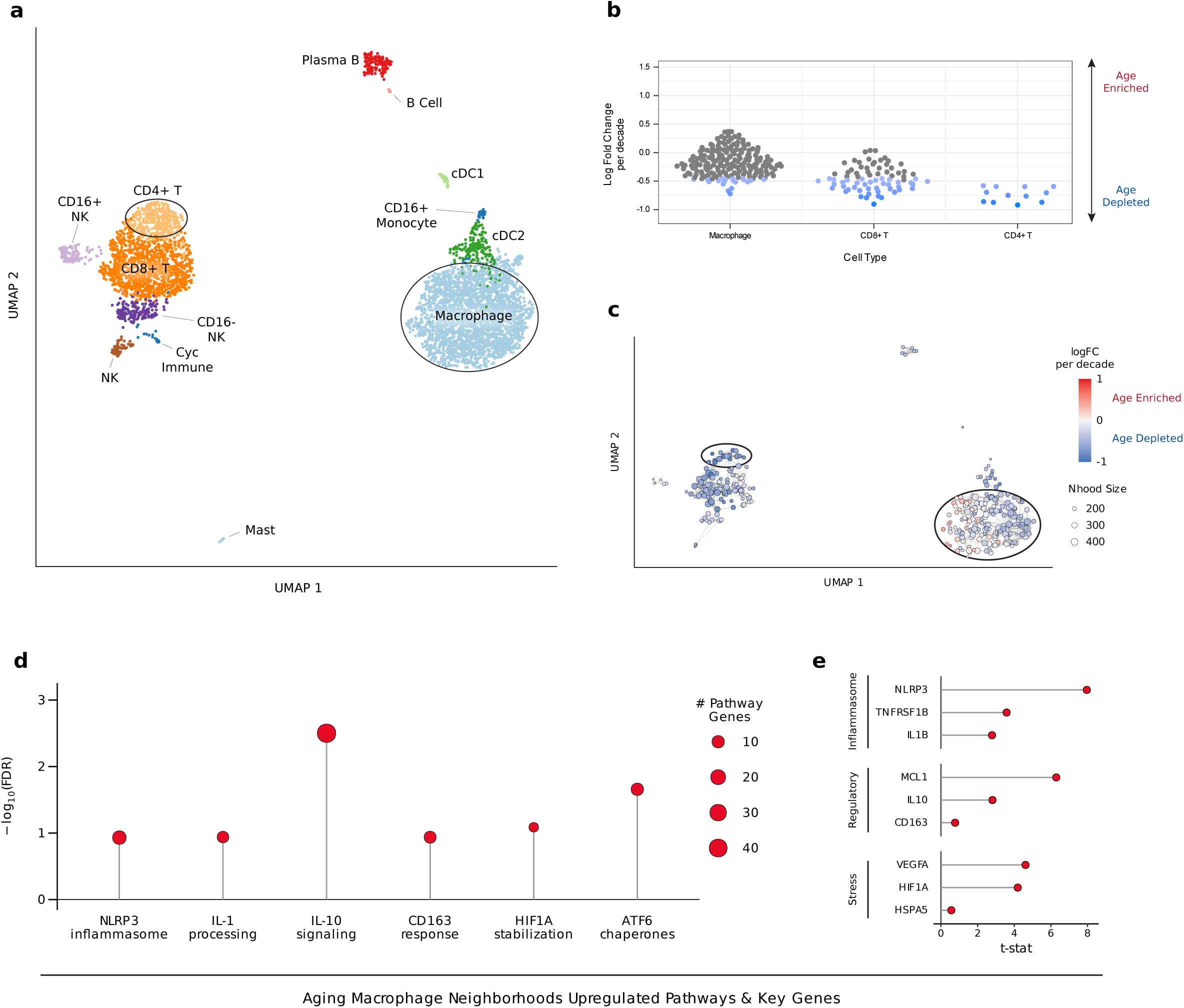
Age-associated remodeling of immune neighborhoods. a, Annotated UMAP of the immune compartment for reference. Key cell types of interest are circled. **b,** Beeswarm plot showing change in Milo neighborhood abundance with age. Neighborhoods with a spatial FDR < 0.1 are colored by direction of change. Cell types represented by fewer than 10 neighborhoods, and neighborhoods with <70% assignment to their dominant cell type annotation, were excluded from plotting. **c,** Milo neighborhood differential abundance effect sizes projected into UMAP space for age. Neighborhoods are colored by log fold change. Key cell types of interest are circled. **d,** Lollipop plot of curated Reactome pathways upregulated in age-enriched relative to age-depleted macrophage neighborhoods. Dot size represents the number of pathway genes included. **e,** Lollipop plot of moderated t-statistics for selected genes grouped into coherent biological themes from the differential expression comparison between age-enriched and age-depleted macrophage neighborhoods. Nhood, neighborhood.

To define the programs distinguishing age-enriched versus age-depleted macrophage neighborhoods, we performed differential expression and pathway analysis on grouped macrophage neighborhoods classified by Milo age-associated log-fold change (see Methods). Age-enriched macrophage neighborhoods showed enrichment for inflammasome-related, regulatory, and stress-response pathways, including NLRP3 inflammasome, IL-1 processing, IL-10 signaling, CD163 response, HIF1A stabilization, and ATF6 chaperones (**Fig. 4d**). Correspondingly, age-enriched neighborhoods upregulated inflammasome-associated genes (*NLRP3*, *IL1B*, *TNFRSF1B*), regulatory response genes (*IL10*, *CD163*, *MCL1*), and stress-response genes (*HIF1A*, *VEGFA*, *HSPA5*) (**Fig. 4e**). Several of these genes also showed distinct expression gradients across the macrophage region of the UMAP from age-depleted to age-enriched neighborhoods (**Extended Data Fig. 7a**). The coexistence of inflammasome, regulatory, and cellular-stress programs argues against a uniform pro-inflammatory macrophage shift. Instead, it suggests age-associated redistribution among transcriptionally distinct macrophage states.

To test whether macrophage neighborhood-level differential expression reflected signals beyond those detected when macrophages were analyzed as a single annotated cell type in the following section, we compared gene-level t-statistics between the two approaches. Although directionally concordant, the two sets of t-statistics were only modestly correlated (ρ = 0.35), indicating that Milo captured partially distinct intra-macrophage signals rather than simply recapitulating the whole-cell-type age signature at finer scale (**Extended Data Fig. 7b**).

### Moderate parity is associated with compositional and transcriptional patterns opposing ovarian aging

Having characterized age-associated remodeling of ovarian cellular composition and immune states, we next examined whether reproductive history was associated with these same features. Focusing on the primary comparison of moderate parity (1–3 live births) with nulliparity (see Methods), we compared age- and parity-associated effects across cell type abundances, immune neighborhoods, and cell type-specific transcriptional programs.

Moderate parity was associated with a compositional pattern that broadly opposed the age-associated shifts observed in the snucRNA-seq atlas (**Fig. 5a**). Donors with 1–3 live births showed higher relative abundance of CD4+ T cells and lower relative abundance of TNF-activated stromal cells than nulliparous donors, opposing the age-associated depletion and enrichment of these respective populations (**Fig. 5b,c**). Opposing directions were also evident across several other stromal, vascular, and immune populations, although this pattern was not uniform across cell types (**Fig. 5a**). Interestingly, some populations showed nonlinear relationships across increasing parity levels in the raw proportions (Supplementary Fig. 4; Supplementary Note 4). These findings associate reproductive history with persistent differences in postmenopausal ovarian cellular composition that frequently oppose age-associated remodeling.

**Fig 5.**
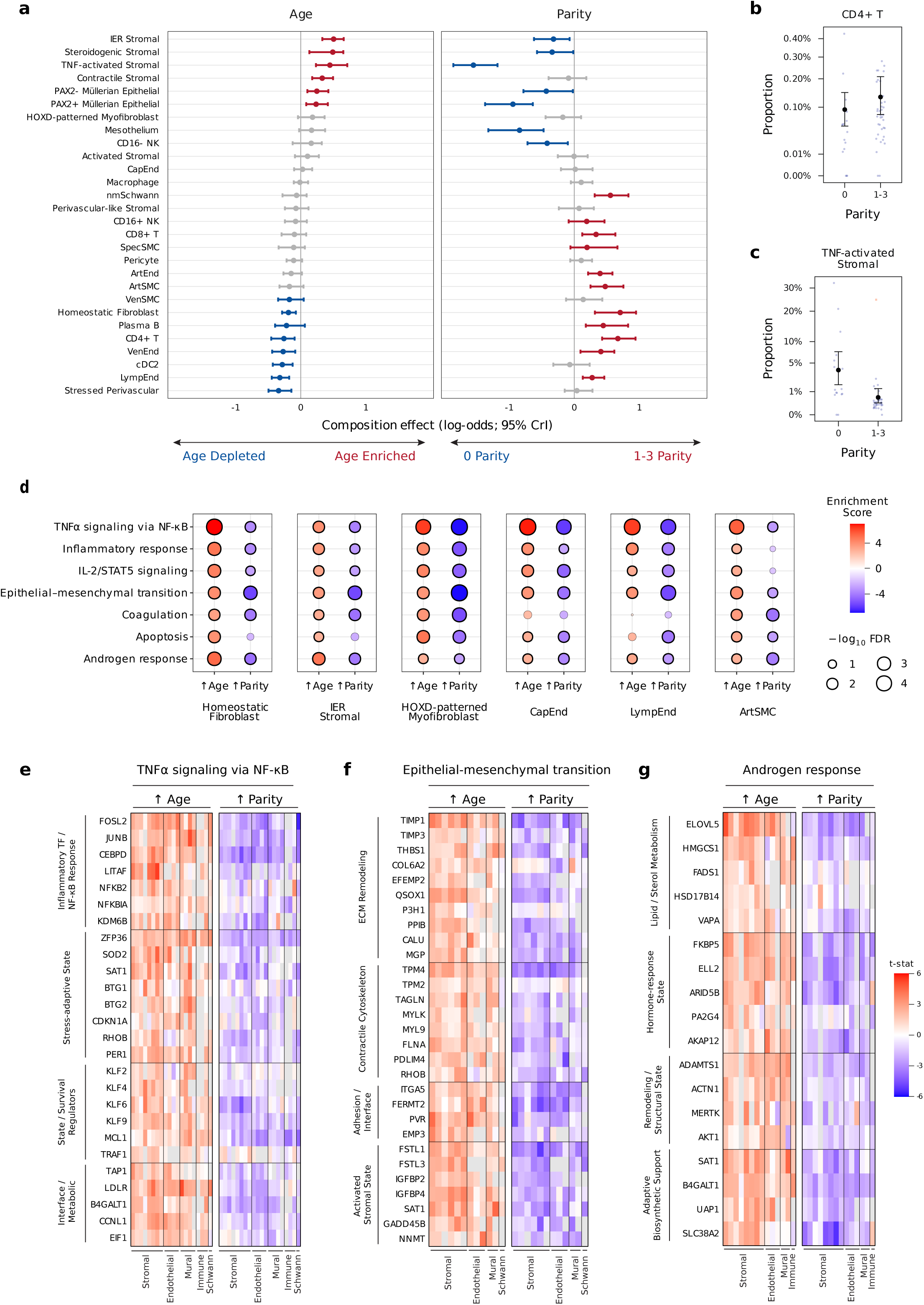
Moderate parity and aging show opposing associations for cell composition and gene expression. a, Forest plots showing age-associated changes in cell type abundance beside parity-associated changes distinguishing nulliparous (0 live births) donors from donors with moderate parity (1-3 live births). Points show posterior median composition effects on the log-odds (logit) scale, expressed per one-standard-deviation increase in age or for moderate parity (1–3 live births) relative to nulliparity (0 live births). Error bars indicate 95% credible intervals. Colored entries indicate significant effects (Bayesian FDR ≤ 0.05) based on posterior evidence for an absolute composition effect exceeding 0.1 logit units. The left aging panel is the same as the left panel from Figure 3a, displayed here as a visual comparator to the parity contrast. **b–c,** Model-predicted proportions for representative cell types across parity groups. Large black points indicate posterior median proportions and error bars indicate 95% credible intervals. Small blue points indicate raw sample proportions, and red squares indicate sample–cell type pairs censored as outliers by sccomp. **d,** Dotplots showing recurrent MSigDB Hallmark pathways changing with age across highlighted cell types, with corresponding parity results shown alongside. Dot color indicates pathway Enrichment Score, defined as the pathway-level t-statistic, and dot size indicates −log10 FDR. **e–g,** Heatmaps showing selected genes contributing to highlighted age-enriched pathways. Genes are arranged in rows and grouped by biological theme; cell types are arranged in columns and grouped by compartment. Colors represent gene-level moderated t-statistics from dreamlet mixed-effects models, with corresponding parity results shown alongside the age results. Upward parity arrows for gene and pathway analyses represent the effect of moderate parity (1-3 live births), as compared to nulliparity (0 live births).

At the neighborhood level, Milo analysis identified parity-associated enrichment of CD8+ T and CD4+ T cell neighborhoods in donors with 1–3 live births relative to nulliparous donors, contrasting with the age-associated depletion observed within these populations (**Extended Data Fig. 8**). These findings extended the opposing compositional patterns to transcriptionally defined neighborhoods within the adaptive immune compartment.

We next asked whether this relationship extended to transcriptional programs within annotated cell types. Using a pseudobulk linear mixed-model framework that accounted for biological and technical covariates, we estimated age- and parity-associated gene expression changes (Supplementary Table 6) and summarized the top moderated t-statistics within each cell type using a root-mean-square perturbation score (see Methods). Stromal and vascular-associated cell types exhibited some of the strongest transcriptional changes associated with both variables (**Extended Data Fig. 9a**). Notably, capillary endothelial cells were the most transcriptionally perturbed with age despite exhibiting minimal changes in relative abundance, highlighting a distinction between cell-intrinsic transcriptional remodeling and shifts in cell type abundance. This finding indicates that the ovarian microvasculature shows pronounced age-associated transcriptional differences even in the absence of a detectable change in capillary endothelial cell abundance.

To identify higher-order biological programs associated with age and parity, we next performed Hallmark pathway analysis across cell types (see Methods). Rather than being restricted to individual cell types, several pathways were repeatedly perturbed across stromal, endothelial, and mural populations, including TNFα signaling via NF-κB, epithelial–mesenchymal transition, apoptosis, and androgen response (**Fig. 5d**; **Extended Data Fig. 9b**). Across several cell types, pathways enriched with increasing age showed opposing enrichment patterns in donors with moderate parity relative to nulliparous donors, consistent with the age–parity opposition observed in cellular composition.

To examine the gene-level patterns underlying these pathway results, we focused on three pathways recurrently enriched across cell types: TNFα signaling via NF-κB, epithelial–mesenchymal transition, and androgen response. Within these pathways, we identified genes showing concordant age-associated increases across cell types and compared their corresponding parity-associated effects (**Fig. 5e–g**). Across all three pathways, the parity-associated effects for these same genes frequently opposed the age-associated direction, extending the age–parity opposition from pathway-level enrichment to shared gene-level programs.

For TNFα signaling via NF-κB, concordant age-associated genes across cell types included canonical inflammatory response genes such as *NFKBIA*, *JUNB*, and *CEBPD*, the latter of which has recently been implicated in human ovarian aging^6^. Additional shared genes reflected a broader stress-adaptive state, including *ZFP36*, *SOD2*, and *CDKN1A*, together with KLF-family, *MCL1*, and *TRAF1* survival-associated regulators and metabolic/interface genes such as *LDLR* and *B4GALT1* (**Fig. 5e**). Collectively, these transcriptional patterns are consistent with a fibro-inflammatory and stress-adaptive state that shares features with senescence-associated TNFα/NF-κB signaling programs described in aging tissues^49–52^.

For epithelial–mesenchymal transition, the top shared age-associated genes did not primarily indicate epithelial-cell transdifferentiation but instead reflected a broader stromal remodeling program captured by this Hallmark pathway. Shared genes spanned extracellular-matrix remodeling (*THBS1*, *TIMP1*, *COL6A2*), contractile cytoskeletal activation (*TAGLN*, *MYLK*, *MYL9*), adhesion/interface biology (*ITGA5*, *FERMT2*), and an activated stromal-state module (*FSTL1*, *IGFBP2*, *NNMT*) (**Fig. 5f**). These patterns are consistent with fibrosis-like remodeling and contractile stromal activation, in line with prior studies describing age-associated extracellular-matrix and stromal remodeling in the ovary^7,14^.

For androgen response, shared age-associated genes were dominated by a hormone-responsive lipid and sterol metabolic program, including *HMGCS1*, *FADS1*, *ELOVL5*, and *HSD17B14*, together with remodeling and adaptive support genes such as *ADAMTS1*, *MERTK*, *B4GALT1*, and *SLC38A2* (**Fig. 5g**). In the context of the age-associated expansion of steroidogenic stromal cells, these patterns support an age-associated shift toward steroidogenic and androgen-responsive biology in the postmenopausal ovary.

Together, these analyses show that moderate parity was associated with both differences in cellular composition and cell type transcriptional programs that frequently opposed the corresponding age-associated patterns. This relationship spanned several ovarian compartments, encompassing immune populations, stromal remodeling, and inflammatory and hormone-responsive programs.

### Cell–cell communication highlights stromal-macrophage interaction with aging and parity

To assess whether age- and parity-associated remodeling was also reflected in inferred multicellular communication programs, we analyzed putative ligand–receptor interactions within each sample using LIANA+^53^ and decomposed the resulting communication tensor with Tensor-cell2cell^54^ (see Methods). This analysis identified 10 inferred communication programs with distinct cellular and ligand–receptor interaction patterns (**Extended Data Fig. 10**; Supplementary Table 7). These encompassed mesenchymal–endothelial matrix interactions, perivascular-to-stromal extracellular-matrix remodeling, and stromal-to-mural remodeling (Factors 1–3); lymphocyte recruitment (Factor 4); endothelial remodeling and reciprocal endothelial signaling (Factors 5–6); stromal–macrophage scavenging-associated communication (Factor 7); reciprocal mural support and endothelial-to-mural NOTCH signaling (Factors 8–9); and macrophage-dominant outgoing communication (Factor 10) (**Extended Data Fig. 10a– j,m**).

Among these programs, Factor 7 linked stromal senders to macrophage receivers and was distinguished by APOE–LRP1, SERPINE2–LRP1, and SERPINF1–PLXDC2 interactions, consistent with a putative scavenging- and remodeling-associated communication program (**Extended Data Fig. 10j,m**). At the sample level, Factor 7 scores tended to increase with age and were lower in donors with moderate parity (1–3 live births) than in nulliparous donors, although neither association reached significance after multiple-testing correction (age p_adj_ = 0.12; parity p_adj_ = 0.07; **Extended Data Fig. 10k,l**). These directional patterns were consistent with the broader age–parity opposition observed in compositional and transcriptional analyses. Together, these findings identify a putative stromal–macrophage communication program whose directional associations with age and parity align with the broader remodeling observed across the postmenopausal ovary, providing a candidate multicellular axis for future investigations.

## Discussion

Here, we generated, to our knowledge, the largest reported cell-type-resolved atlas of the aging human ovary from a single donor cohort, spanning 28 unique postmenopausal donors aged 50–84 years, and resolved 37 fine cell states. Integrating compositional, cell-intrinsic transcriptional, neighborhood-level, and inferred cell–cell communication analyses, with key compositional findings further supported in an external cohort, we found that the postmenopausal ovary is not an inert follicle-depleted remnant, but instead a biologically active aging tissue exhibiting stromal–immune remodeling, vascular loss, and a shift toward steroidogenic and androgen-responsive biology. We further identified broad opposition between age-associated signatures and those associated with moderate parity at both molecular and compositional levels, consistent with obstetric history leaving a detectable molecular and cellular signature in the postmenopausal ovary.

Our data suggest a nuanced view of inflammatory remodeling in the postmenopausal ovary, characterized not by simple immune cell accumulation but by reduction of immune populations together with expansion of stress-associated and TNF-activated stromal states. Age-associated reductions in T cell populations were directionally concordant between the snucRNA-seq atlas and the external GTEx cohort. Within the snucRNA-seq atlas, homeostatic fibroblasts declined in relative abundance, whereas immediate-early response and TNF-activated stromal states increased. This compositional pattern was paralleled by cell-intrinsic transcriptional changes, with recurrent enrichment of TNFα signaling via NF-κB across stromal and vascular-associated cell types and concordant upregulation of inflammatory and stress-adaptive genes including *NFKBIA*, *JUNB*, *CEBPD*, *ZFP36*, *SOD2*, and *CDKN1A*, consistent with prior human and murine data implicating NF-κB signaling, *CEBPD*, and *CDKN1A* in ovarian aging^6,7,22^. Collectively, these features are also compatible with chronic fibro-inflammatory and senescence-associated stress signaling programs described in other aging tissues^49–52^. Prior human and mouse studies likewise support stromal remodeling and immune reorganization during ovarian aging; however, several murine studies reported increased lymphoid abundance with age^22,55–57^, whereas recent human comparisons of reproductively young and aged ovaries did not detect major immune compositional differences^6^. Because this study was cross-sectional and restricted to postmenopausal tissue, these associations cannot establish longitudinal aging trajectories or determine when the changes emerge across the reproductive aging continuum. In this context, our findings, which span multiple decades of postmenopausal aging, suggest that immune remodeling in the postmenopausal ovary may differ from reproductive aging and may involve age-associated reductions in lymphoid representation that could diminish immune surveillance and coordination, potentially permitting persistence of chronic stress-associated stromal states.

Neighborhood-level Milo analysis provided complementary support for the immune compositional changes identified by cell type modeling, while also revealing heterogeneity not captured by aggregate cell type abundances. In particular, age-associated depletion of CD4+ and CD8+ T cell neighborhoods mirrored the directional changes observed in the cell type compositional analysis despite the use of a distinct neighborhood-graph framework, supporting the robustness of these lymphoid declines. By contrast, macrophage neighborhoods showed a wider spectrum of age-associated effect sizes, ranging from significant depletion to modest positive shifts across the cluster. Macrophage neighborhoods with positive age-associated log-fold changes, compared with those with negative age-associated log-fold changes, were characterized by genes and pathways related to the NLRP3 inflammasome, regulatory macrophage programs, and cellular stress responses, consistent with murine ovarian aging studies implicating NLRP3 inflammasome activation^23^, broader aging literature linking NLRP3 signaling to macrophage aging and immune homeostasis^58^, and human ovarian aging data from reproductive-age donors showing enrichment of pyroptotic macrophages in middle-aged relative to young ovaries^11^. Importantly, the coexistence of inflammasome, regulatory, and stress programs among macrophage neighborhoods with positive age-associated shifts argues against a uniform pro-inflammatory response and instead supports state-specific macrophage remodeling with age.

Aging was also associated with marked vascular remodeling, particularly involving depletion of venous and lymphatic endothelial populations together with loss of arterial and venous smooth muscle support, whereas capillary endothelial cells showed relatively limited compositional change yet emerged as the most transcriptionally perturbed vascular population. This distinction suggests that vascular aging in the postmenopausal ovary is not fully captured by abundance shifts alone but also involves substantial remodeling within the residual microvasculature. Consistent with this, endothelial populations upregulated genes associated with epithelial–mesenchymal transition and related remodeling programs, with shared genes highlighting extracellular-matrix remodeling, adhesion, and contractile activation. Prior human and murine studies likewise support vascular involvement in ovarian aging, with human atlases reporting age-associated shifts in vascular cell abundance^6^ and recent work in mice suggesting that ovarian vascular decline may be a primary driver of ovarian aging rather than merely a downstream consequence^24^. In the context of the well-described shrinkage and atrophy of the postmenopausal ovary^59^, these findings raise the possibility of disproportionate remodeling of the vascularized medullary compartment, highlighting the value of cortex-medulla resolved profiling as previously demonstrated in postmenopausal ovarian tissue^60^. The pronounced transcriptional perturbation of capillary endothelium further suggests continued remodeling within the remaining microvasculature. Recent murine work demonstrated that impaired VEGFR3-dependent ovarian lymphangiogenesis leads to fibrosis and premature ovarian aging^61^, supporting a potential contribution of lymphatic insufficiency to ovarian remodeling. Although the functional consequences of declining lymphatic representation in the human postmenopausal ovary remain to be established, reduced lymphatic support could plausibly impair tissue clearance and inflammatory resolution^62,63^, potentially contributing to a microenvironment that favors persistent tissue stress and fibrosis.

Our data also support an age-associated shift toward steroidogenic and androgen-responsive biology in the postmenopausal ovary. With age, the ovary showed expansion of steroidogenic stromal cells together with enrichment of androgen response programs across multiple stromal, endothelial, and mural cell types. At the cell-state level, the steroidogenic stromal population was marked by cholesterol biosynthesis and steroid hormone metabolism genes, whereas the contractile stromal state showed theca externa-like features and closely tracked steroidogenic stromal abundance across cohorts. At the transcriptional level, age-associated androgen response genes were dominated by a hormone-responsive lipid and sterol metabolic program, including *HMGCS1*, *FADS1*, *ELOVL5*, and *HSD17B14*. Although our study did not measure circulating hormones or directly assess steroid production, prior human ovarian vein sampling and oophorectomy studies indicate that the postmenopausal ovary can remain an androgen-producing organ capable of contributing to serum testosterone levels^31^. Furthermore, recent mass spectrometry-based population studies showing a late postmenopausal rise in total testosterone raise the possibility that persistent or increasing ovarian steroidogenic activity may contribute to this pattern^32,33^. Together, these results provide a cellular and transcriptional framework for continued steroidogenic activity in the follicle-depleted postmenopausal ovary and suggest that these programs may shape a broader androgen-responsive tissue environment.

A prominent theme of this study was the broad opposition between age- and parity-associated transcriptional and compositional signatures. In the postmenopausal ovary, age-associated loss of large-vessel and lymphatic-associated populations and the shift from homeostatic to stress-associated stromal states were broadly opposed in donors with 1–3 live births relative to nulliparous donors. This opposition was most evident in the increased abundance of CD4+ T cells and reduced abundance of TNF-activated stromal cells with moderate parity, but it also extended to shared transcriptional programs, as inflammatory and tissue-remodeling pathways enriched with age, including TNFα signaling via NF-κB, inflammatory response, and epithelial– mesenchymal transition, were depleted in donors with 1–3 live births. A similar directional pattern was observed for the inferred stromal–macrophage communication program represented by Factor 7, although its age and parity associations did not reach significance after multiple-testing correction.

The detection of these differences decades after reproductive exposure is notable and suggests that reproductive history may leave long-lasting tissue-level signatures associated with several core aging-related processes, including inflammation and intercellular communication^64^. This interpretation is broadly consistent with epidemiological studies linking moderate parity to delayed natural menopause^34^, lower blood metabolic and proteomic biological age^35,65^, and more favorable late life health and survival outcomes^66^, while also suggesting nonlinearity across parity levels^35,36^. Mechanistically, these opposing age- and parity-associated patterns could plausibly reflect persistent immune reprogramming after pregnancy^38,67^, systemic consequences of reproductive history^37,68^ and reduced lifetime ovulatory cycles^69,70^, with pregnancy-associated microchimerism representing an additional, more speculative possibility^71^. However, because parity was defined by live birth count and detailed reproductive and life-course histories, including age at childbirth, breastfeeding, and hormone use, were unavailable, the contributions of pregnancy itself versus correlated exposures cannot be disentangled. More broadly, these findings raise the possibility that moderate parity-associated pathways may inform not only ovarian biology but aging biology more generally, with the postmenopausal ovary providing a tractable tissue context in which to investigate associations between reproductive history and later-life molecular state.

Some additional considerations qualify the interpretation and generalizability of these findings. Our cohort comprised women undergoing surgery for clinical indications and was predominantly White, with limited representation of Black and higher-parity donors, constraining generalizability and subgroup inference. Unmeasured clinical factors may also influence the observed associations. Moreover, compositional estimates from snucRNA-seq reflect relative nuclear recovery rather than absolute cell abundance and may be influenced by cell type-specific differences in nuclear isolation or capture.

Together, these findings position the postmenopausal ovary as an actively remodeled aging tissue rather than a passive follicle-depleted remnant. By integrating compositional, transcriptional, neighborhood-level, and cell–cell communication analyses, our study reveals coordinated stromal, vascular, immune, and steroidogenic remodeling with age, while identifying opposing patterns associated with moderate parity. Future studies that integrate spatial and orthogonal molecular profiling with richer reproductive, endocrine, and clinical metadata will be important for defining how these programs are organized within ovarian microanatomy and how they relate to systemic aging and health outcomes. More broadly, our findings establish the postmenopausal ovary as a valuable system for investigating how reproductive history intersects with molecular and cellular aging later in life.

## Extended Data Figure legends

**Extended Data Fig 1. Ovary donor cohort metadata characterization. a,** Donor metadata covariate association matrix. Pairwise associations were quantified according to variable type using Spearman’s ρ (numeric–numeric), point-biserial correlation (binary–numeric), η² (categorical–numeric), Cramér’s V with Bergsma– Wicher correction (categorical–categorical), or signed φ with Bergsma–Wicher correction (binary–binary). Selected pairwise comparisons between derived or overlapping metadata variables were masked and denoted by “-”. **b–d**, Boxplots with overlaid donor data points showing parity in relation to urinary incontinence, pelvic organ prolapse, and uterine malignancy status. Effect sizes are shown as point-biserial correlation coefficients. P values were calculated using two-sided Mann–Whitney U tests and adjusted using the Benjamini–Hochberg method across the three comparisons. Boxes indicate the interquartile range, center lines indicate medians, and whiskers extend to the most extreme observations within 1.5 times the interquartile range of the corresponding quartile. Overlaid points represent individual donors.

**Extended Data Fig 2. Single-nucleus RNA-seq atlas cell type annotation and integration. a,** Dotplot showing coarse annotated cell types from Fig. 1 and corresponding marker genes. Gene expression was scaled across cell types, and dot size indicates the fraction of nuclei in each cell type with detectable expression. Cell type and gene labels are colored according to broad compartment classification for ease of visualization. **b–d**, UMAP plots with nuclei colored by donor of origin, non-ovarian cancer state, or processing batch.

**Extended Data Fig 3. Annotation of non-stromal cellular compartments. a–b, e–g,** UMAPs showing fine cell type annotations for endothelial, mural, immune, epithelial, and Schwann compartments in the postmenopausal ovary. **c–d, h–j,** Dotplots showing corresponding marker genes for fine annotated endothelial, mural, immune, epithelial, and Schwann cell types. Gene expression was scaled across cell types, and dot size indicates the fraction of nuclei in each cell type with detectable expression. ArtSMC, arterial smooth muscle; VenSMC, venous smooth muscle; SpecSMC, specialized smooth muscle; cDC, conventional dendritic cell; mSchwann, myelinating Schwann; nmSchwann, non-myelinating Schwann.

**Extended Data Fig 4. Characterization of specialized cell states. a,** Distribution of donor-level BuckSenOvary signature scores across cell types. Each point represents the median score across nuclei from one donor within a given cell type. Boxes indicate the interquartile range across donor-level scores, center lines indicate medians, and whiskers extend to the most extreme observations within 1.5 times the interquartile range of the corresponding quartile. **b,** UMAP of the annotated endothelial compartment highlighting a specialized endothelial population. **c,** UMAPs colored by log-normalized expression of prominent marker genes for the specialized endothelial population. **d,** UMAP of the annotated immune compartment highlighting a specialized GPNMB+ macrophage population. **e,** UMAPs colored by log-normalized expression of prominent marker genes for the GPNMB+ macrophage population. SpecEnd, specialized endothelial; Mac, macrophage.

**Extended Data Fig 5. Deconvolved bulk proportion and expression. a,** Stacked bar plot showing deconvolved GTEx ovary cell type proportions. Cell types were grouped into broad compartment categories, and donors were ordered by Homeostatic Fibroblast proportion. A schematic interpretation of putative cortex-enriched to medulla-enriched composition is shown from high to low Homeostatic Fibroblast proportion, respectively. Each bar represents one donor. Illustration was created using BioRender.com. **b,** Pearson correlation matrix of deconvolved cell type gene expression profiles.

**Extended Data Fig 6. Race-associated and co-abundance compositional analysis. a,** Forest plots showing race-associated differences in cell type abundance in the snucRNA-seq atlas and deconvolved bulk GTEx RNA-seq data. Cell types not represented in one of the two analyses were omitted from that panel for clarity. Points show posterior median composition effects on the log-odds (logit) scale with 95% credible intervals. Colored entries indicate significant effects (Bayesian FDR ≤ 0.05) based on posterior evidence for an absolute composition effect exceeding 0.1 logit units. **b,** Model-predicted proportions for representative cell types across race groups. Large black points indicate posterior median proportions and error bars indicate 95% credible intervals. Small blue points indicate raw sample proportions, and red squares indicate sample–cell type pairs censored as outliers by sccomp. **c,** Scatter plots of centered log-ratio (CLR) transformed cell type proportions across all samples in snucRNA-seq and deconvolved bulk datasets. Spearman’s ρ and Benjamini–Hochberg-adjusted P values across all tested cell type pairs within each technology are shown.

**Extended Data Fig 7. Age-associated macrophage expression patterns and methods comparison. a,** Feature UMAPs of the immune compartment colored by log-normalized expression of selected marker genes identified from the differential expression comparison between age-enriched and age-depleted macrophage neighborhoods. **b,** Scatter plot comparing gene-level t-statistics from the macrophage compartment in the dreamlet linear mixed-effects framework and the differential expression results comparing Milo age-enriched versus age-depleted macrophage neighborhoods. Each dot represents one gene. Spearman’s ρ is shown.

**Extended Data Fig 8. Immune neighborhood differential abundance with age and parity. a–b,** Beeswarm plots showing change in Milo neighborhood abundance with age or parity group. Neighborhoods with a spatial FDR < 0.1 are colored by direction of change. Cell types represented by fewer than 10 neighborhoods, and neighborhoods with <70% assignment to their dominant cell type annotation, were excluded from plotting. **c**, Annotated UMAP of the immune compartment for reference. Key cell types of interest are circled. **d–e,** Milo neighborhood differential abundance effect sizes projected into UMAP space for age or parity group. Neighborhoods are colored by log fold change. Key cell types of interest are circled. Age effects shown are per decade. Positive parity log-fold changes indicate enrichment in donors with 1–3 live births relative to donors with 0 live births. Nhood, neighborhood.

**Extended Data Fig 9. Gene expression and pathway analysis across all tested cell types. a,** Lollipop plots showing all tested cell types ranked by transcriptional perturbation for age and parity (0 versus 1–3 live births), calculated as the root-mean-square of the top 500 genes by absolute moderated t-statistic from dreamlet mixed-effects models. Perturbation scores are direction-agnostic and were min–max scaled from 0 to 1 across cell types. **b,** Dotplots showing MSigDB Hallmark pathways changing with age across all tested cell types, with corresponding parity results shown alongside. Displayed pathways comprise a non-redundant set of the top 3 significant pathways by absolute t-statistic for each cell type and contrast. Dot color indicates pathway Enrichment Score, and dot size indicates −log10 FDR; dots with FDR ≤ 0.05 are shown with a thicker outline.

**Extended Data Fig 10. Cell–cell communication characterization. a–j,** Factor network plots showing inferred cell–cell communication programs between sending and receiving cell types for latent communication factors derived from LIANA+ and Tensor-cell2cell. Cell type nodes are colored by broad compartment for ease of visualization. **k–l,** Associations between Factor 7 score and donor age or parity group from ordinary least-squares models with donor-clustered standard errors. In **k**, the line and ribbon show the fitted relationship and 95% confidence interval; in **l**, the large dot and error bars show the model-estimated mean and 95% confidence interval. Small dots represent individual samples. Displayed P values were adjusted using the Benjamini– Hochberg method across all factors within each model term. **m,** Curated ligand–receptor (LR) pairs distinguishing all latent communication factors. Non-redundant and biologically coherent LR pairs were selected for display. Heatmap colors represent LR-pair loadings per factor, z-scored across factors.

## Methods

### Experimental Methods

#### Tissue acquisition and handling

De-identified human ovarian tissue was obtained from donors through the Northwestern University Reproductive Tissue Library (NU-RTL) under Institutional Review Board-approved protocols (STU00215770 and STU00215938). Ovarian tissue was collected from women aged 50–84 years undergoing bilateral salpingo-oophorectomy and/or total laparoscopic hysterectomy for benign gynecologic conditions, endometrial malignancy or premalignancy, or cervical premalignancy (Supplementary Table 1). Only cases without ovarian neoplasia were included, and donors with endometriosis were excluded.

Upon collection, ovaries were sectioned into 3–5 mm thick cross-sections perpendicular to the long axis of the ovary. In the absence of significant gross pathology, as assessed by a certified gynecologic pathologist, up to two cross-sections were designated for research and transported on ice in ORIGIO Handling IVF medium (CooperSurgical, Trumbull, CT, USA). One half of each cross-section was subdivided in a 60 mm glass Petri dish into 20–30 mg tissue pieces containing both cortex and medulla. Tissue pieces were washed in PBS, flash-frozen in 1.5 mL tubes, shipped overnight to the Buck Institute for Research on Aging on dry ice, and stored at −80°C upon arrival.

#### Single-nucleus RNA-seq library preparation and sequencing

Cryopreserved ovarian tissue pieces containing both cortex and medulla were processed for fixed tissue dissociation using the 10x Genomics Chromium Fixed RNA Profiling tissue fixation and dissociation protocol (CG000553 Rev B), with protocol-specific optimization for human ovary tissue. Fixed tissue was dissociated in RPMI containing 0.2 mg ml⁻¹ Liberase TH using a gentleMACS Octo Dissociator with Heaters, and two additional “spin only” cycles were performed to further disrupt residual large tissue fragments. Fixed nuclei suspensions were filtered through 30 μm strainers and counted using a Cellaca PLX automated cell counter with AO/PI staining.

Samples were then processed using the 10x Genomics Chromium Fixed RNA Profiling Reagent Kits for Multiplexed Samples (CG000527 Rev E). Fixed suspensions were hybridized overnight with uniquely barcoded whole-transcriptome probe sets, pooled after hybridization, washed, and loaded onto a Chromium Next GEM Chip Q for GEM generation and barcoding targeting 8,000 nuclei per sample. Following this, post-GEM recovery, pre-amplification, and sample indexing PCR steps were performed according to the manufacturer’s protocol. Samples were multiplexed in batches of 16 using unique probe barcodes; across the study, 64 ovarian tissue pieces from 28 donors were processed across 4 plexes (Supplementary Fig. 1). Libraries were assessed on an Agilent 4200 TapeStation for size distribution and concentration, then sequenced on an Element Biosciences Aviti instrument with 10% PhiX spike-in. Sequencing was performed with the following read configuration: Read 1, 29 bp; Read 2, 91 bp; Index 1, 10 bp; and Index 2, 10 bp, yielding an average of ∼14,000 read pairs per nucleus.

Sample processing logs are available in Supplementary Table 1 and expanded protocol details are available on https://www.protocols.io/ ^72^.

### Computational Methods

#### snucRNA-seq preprocessing, integration, and annotation

Sequencing data were processed with the 10x Genomics Cell Ranger *multi* pipeline (v8.0.0) using the human GRCh38-2024-A reference and the human Flex probe set (v1.0.1), generating per-sample raw and filtered feature-barcode matrices. Ambient RNA was reduced on a per-sample basis with CellBender^73^ *remove-background* (v0.3.2) applied to the raw matrices, and corrected matrices were subsequently restricted to cell barcodes retained in the corresponding Cell Ranger filtered matrices. Downstream analyses were performed using Python (v3.9 and v3.12) and R (v4.4).

Quality-control metrics, including mitochondrial transcript percentage, were calculated with scanpy^74^ *pp.calculate_qc_metrics* (v1.10.2). Initial doublet calls were generated with scDblFinder^75^ (v1.18.0). Putative low-quality nuclei were then removed if they exceeded 5 median absolute deviations (MADs) from the median for total counts or detected genes, or if mitochondrial transcript percentage exceeded 10%.

For integration, the top 10,000 highly variable genes were selected with scanpy *pp.highly_variable_genes* using the Seurat^76^ v3 flavor, and a latent representation was learned with scVI from scvi-tools^77^ (v1.1.6). Unique sample ID was used as the batch variable, donor ID and processing batch were included as categorical covariates, and mitochondrial transcript percentage was included as a continuous covariate.

Doublets were then further identified using Solo^78^, a doublet-classification framework built on the trained scVI model, followed by high-resolution Leiden clustering. Clusters were removed if their median doublet scores from both Solo and scDblFinder exceeded 0.15. Additional putative doublet clusters were manually identified and removed during compartment-level annotation based on mixed expression of cell type-specific marker genes.

After doublet removal, scVI integration and Leiden clustering were repeated and a putative ambient RNA-enriched cluster was removed. Cell-cycle scores were calculated using scanpy *tl.score_genes_cell_cycle* with previously published marker sets^79^. Major cell compartments were identified using canonical marker genes, subset from the full atlas, and reprocessed independently using the same scVI integration, dimensional reduction, and clustering workflow used for the full dataset. Cell type and cell state annotations within each compartment were assigned using canonical markers from prior literature, *scvi.model.SCVI.differential_expression*, scanpy *tl.rank_genes_groups* on log-normalized counts with the Wilcoxon rank-sum test, and CellTypist^80,81^ (v1.7.1), where an appropriate reference model was available. Annotated compartments were then recombined to generate the final atlas for visualization and downstream differential expression, pathway, compositional, and cell–cell communication analyses.

#### Derivation of sample-level StressScore

To account for transcriptional stress responses that arise during tissue collection prior to freezing and fixation, we derived a sample-level StressScore from a previously reported immediate-early response signature showing cross-dataset concordance across multiple tissues and agreement with both murine dissociation stress and human postmortem tissue (Supplementary Note 2)^82^. Per-nucleus signature scores were computed with PyUCell^83,84^ *compute_ucell_scores* (v0.4.0) and robustly z-scored within each cell type. To obtain a composition-robust sample-level covariate, per-nucleus scores were aggregated by sample and cell type during pseudobulk construction, and StressScore was defined for each sample as the median across cell types of the aggregated mean z-scored signature value. StressScore showed no strong correlation with donor age (Supplementary Fig. 2), supporting its inclusion as a covariate for collection-associated transcriptional stress. StressScore was included as a covariate in downstream analyses where indicated.

#### Sample metadata and modeling conventions

Throughout downstream analyses, snucRNA-seq analyses were performed at the individual tissue piece/sample (unique_sample_ID) level, while Donor ID (SenNet_ID) denoted the donor from whom one or more samples were obtained. Age was modeled as a continuous variable in years unless otherwise specified. Parity was grouped as 0, 1–3, or 4+ live births, with the primary contrast of interest defined as 0 versus 1–3 live births; this grouping was used because the 4+ group was sparse and exploratory analyses suggested nonlinearity in trends across parity levels (see Supplementary Note 4). Processing Batch corresponded to the snucRNA-seq processing batch (Plex) and sparse race categories were collapsed where necessary for stable inference, yielding White, Black, and Other in snucRNA-seq analyses. Repeated sampling from the same donor was accounted for using random effects, donor-blocking, or donor-clustered standard errors where appropriate.

Diagnosis of non-ovarian malignancy was evaluated as a potential covariate but was not included in parity-focused models. In this cohort, most non-benign cases involved endometrial malignancy, and parity was negatively associated with uterine malignancy (**Extended Data Fig. 1**). We therefore prioritized parity as the more biologically proximal and more granular variable for modeling reproductive-history-associated biology, whereas diagnosis of non-ovarian malignancy was treated as a broader clinical summary variable that could overlap with the same etiologic axis and may also reflect local or systemic field effects.

#### Differential gene expression (DGE) analysis

DGE was performed using a pseudobulk linear mixed-model framework implemented in dreamlet^85^ (v1.4.1). CellBender-corrected counts were summed for each annotated cell type within each sample (tissue piece) to generate pseudobulk profiles using *aggregateToPseudoBulk*.

Pseudobulked data were processed with *processAssays* and analyzed with *dreamlet*. Cell types were retained if they contained at least 5 nuclei per pseudobulk sample in at least 10 samples. Within each retained cell type, genes were required to have at least 5 counts in at least 40% of samples (*dreamlet* default). Gene-level models were then fit separately for each cell type using robust empirical Bayes shrinkage under the linear mixed model:

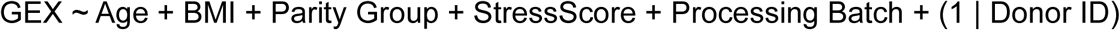

where GEX denotes gene expression for a given gene within a given cell type. P values were adjusted using the Benjamini–Hochberg (BH) method separately for each cell type–predictor combination.

To descriptively rank cell types by transcriptional perturbation for Age or Parity (0 vs 1– 3), genes within each cell type were ranked by the absolute value of their moderated t-statistics and the top 500 genes were retained. Cell type ranking was robust to the number of top genes selected (Supplementary Fig. 10). A perturbation score was then calculated as the root-mean-square (RMS) of the corresponding moderated t-statistics, providing a direction-agnostic summary of transcriptional perturbation strength. For visualization, RMS scores were min–max scaled from 0 to 1 across cell types separately for each contrast.

#### Bulk RNA-seq (GTEx) deconvolution

To assess the generalizability of findings from the postmenopausal human ovary snucRNA-seq atlas, human ovary bulk RNA-seq data and donor metadata were obtained from GTEx^47^ v10 and deconvolved using the annotated snucRNA-seq atlas as a reference. To enrich for likely postmenopausal ovaries, samples were restricted to donors with either an explicit postmenopausal histology annotation and age ≥40 years, or donor age ≥55 years. Samples from sparse race and tissue collection center categories were excluded.

To prepare the snucRNA-seq reference, several cell states that were donor-specific, weakly distinguishable in mixed bulk samples, or that degraded deconvolution performance were excluded. Because erythrocyte contamination was evident in GTEx bulk RNA-seq but not captured by the ovary snucRNA-seq atlas, an erythrocyte reference was added from the Tabula Sapiens^12^ – Immune dataset. Erythrocyte-annotated cells were subset from Tabula Sapiens and reviewed using scVI, Leiden clustering, marker expression, and CellTypist-assisted annotation, confirming a predominantly late erythroid population. To balance donor contribution, each reference cell type was downsampled to at most 100 nuclei/cells per donor per cell type. The ovary snucRNA-seq and Tabula Sapiens references were then merged after harmonizing gene identifiers and restricting to overlapping genes with the GTEx bulk expression matrix.

Deconvolution was performed with BayesPrism^86^ (v2.2.2) preprocessing followed by InstaPrism^87^ (v0.1.6). Reference genes were filtered using *cleanup.genes* to remove lowly expressed genes (present in < 5 nuclei/cells) and genes prone to low specificity or technical bias, including ribosomal, mitochondrial ribosomal, other ribosomal-related, mitochondrial, *MALAT1*, and sex-chromosome genes, after which only protein-coding genes were retained. A BayesPrism prism object was then constructed using the filtered reference and GTEx bulk count matrix, with cell types specified as cell.type.labels and gene outlier handling set to outlier.cut = 0.01 and outlier.fraction = 0.1. Deconvolution was first run with *InstaPrism*, followed by *InstaPrism_update* to perform a second round of estimation using the updated reference matrix. Estimated cell type proportions were carried forward to downstream cell type compositional analyses.

Deconvolution quality was assessed descriptively by extracting top marker genes for each deconvolved cell type, scoring deconvolved expression profiles against canonical and atlas-derived marker signatures using UCell^84^ (v2.12.0), and examining Pearson correlations among deconvolved cell type expression profiles. Based on this review, deconvolved profiles were relabeled where appropriate, merged when they represented indistinguishable categories, and transcriptionally ambiguous or non-informative profiles were excluded from downstream reporting (see Supplementary Note 3).

#### Cell type compositional analysis with sccomp

To assess compositional changes across annotated cell types in both the snucRNA-seq atlas and the GTEx deconvolved bulk RNA-seq data, we used sccomp^88^ (v2.1.23), a Bayesian compositional modeling framework based on a sum-constrained beta-binomial model that accounts for compositional dependence, outliers, mixed-effects designs, and cell type-specific variability.

For snucRNA-seq, cell types were required to contain at least 100 nuclei and be present in at least 10 donors. Cell types passing filtering were analyzed with *sccomp_estimate* using a bimodal mean–variability association and the default Pathfinder inference algorithm, followed by outlier removal with *sccomp_remove_outliers* and significance testing with *sccomp_test*. Composition was modeled using the formula:

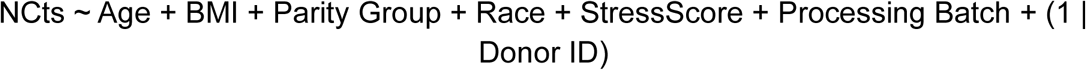

and variability using:

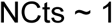

where NCts represents the cell type counts per sample. The variability model ∼ 1 allowed each cell type to have its own baseline variability without fitting covariate-dependent changes in variability.

For deconvolved GTEx bulk RNA-seq proportions, donors with missing values for modeled covariates were removed (104 of the 106 deconvolved samples were retained), sparse Hardy score levels were collapsed, and deconvolved proportions were multiplied by 100,000 and rounded to integers to generate pseudocounts for sccomp (see Supplementary Fig. 7). Cell types with zero counts in more than 50% of donors were excluded. The same sccomp workflow was then applied using the composition formula:

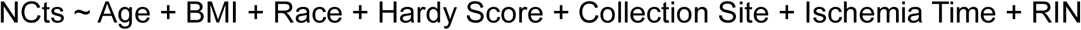

and the variability formula:

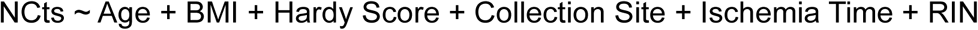

where NCts represents the pseudocounts per cell type per donor, Race represents White (reference) or Black, Hardy Score represents death classification, Collection Site is the tissue collection center, Ischemia Time is sample ischemic time, and RIN is the RNA integrity number. Because race was associated with differences in the abundance of some cell types in both snucRNA-seq and bulk deconvolution analyses, race was included in the composition model but excluded from the variability model to avoid attributing biologically meaningful abundance differences to variability effects.

For both datasets, sccomp internally centered continuous predictors and scaled them to unit standard deviation. Age coefficients therefore represent effects per one-standard-deviation increase in age, with standardization performed across tissue samples for snucRNA-seq and donors for GTEx. Statistical significance was assessed using sccomp’s Bayesian FDR framework, with posterior evidence evaluated against a minimum absolute composition-effect threshold of 0.1 logit units. To improve interpretability, fitted sccomp models were projected back onto the proportion scale using *sccomp_predict* under marginalized nuisance-variable settings. For snucRNA-seq, nuisance variables were marginalized over Donor ID and Processing Batch; for deconvolved bulk data, nuisance variables were marginalized over Hardy Score and Collection Site. The most frequent categorical levels were used as reference levels for marginalization, Age = 55 years was used as the reference age, and the median was used for the remaining numeric variables. Age-associated changes were displayed as fold changes relative to age 55, whereas categorical contrasts were displayed on the proportion scale using marginalized model predictions.

Sensitivity to posterior inference method was evaluated by refitting the primary snucRNA-seq age and parity sccomp models using Hamiltonian Monte Carlo inference. Posterior distributions from these sensitivity analyses are summarized in Supplementary Note 5 and Supplementary Figures 8 & 9.

#### Cell type co-abundance analysis using centered log-ratio (CLR) transformation

To assess cross-sample co-abundance relationships between cell types in both snucRNA-seq and deconvolved bulk RNA-seq data, sample-level cell type proportions were calculated and assembled into sample × cell type proportion matrices. When zero proportions were present, a small pseudocount was added and proportions were renormalized within sample prior to analysis. Proportions were then transformed using the centered log-ratio (CLR) transformation on a per-sample basis to mitigate compositional negative-correlation bias. Pairwise Spearman correlations were calculated across samples for all cell type pairs, and Benjamini–Hochberg correction was applied separately within each technology. For visualization, the relationship between Steroidogenic Stromal and Contractile Stromal proportions was highlighted.

#### Immune neighborhood differential abundance analysis with Milo

To assess variation in immune cell abundance within annotated cell types, we used Milo^48,89^ (v2.6.0), which performs differential abundance testing on partially overlapping neighborhoods defined on a *k*-nearest neighbor graph. Analysis was restricted to the immune compartment, which was of tractable size for neighborhood-level testing and showed notable differences in cell type compositional analyses. Neighborhoods were defined on the immune-compartment scVI latent representation generated during immune-cell annotation using k = 100 nearest neighbors and prop = 0.1 neighborhood seeding.

Prior to differential abundance testing, neighborhoods were required to include nuclei from at least 10 donors and 15 samples (tissue pieces), and the top 3 contributing donors were required to account for no more than 80% of nuclei in a neighborhood, in order to exclude neighborhoods dominated by a small number of donors or samples.

Neighborhood counts per sample were then tested using *testNhoods* with TMM normalization and the HE-NNLS GLMM solver using the design formula:

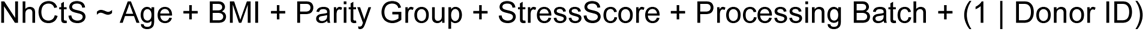

where NhCtS represents the neighborhood counts per sample and Age denotes donor age scaled in 10-year units. For each neighborhood, significance of the coefficient of interest was assessed using a Wald-type test from the mixed-effects model, followed by graph-overlap spatial FDR correction across neighborhoods to account for non-independence of neighborhood membership.^89^

To identify genes distinguishing age-enriched versus age-depleted macrophage neighborhoods, neighborhoods annotated as >99% macrophage were retained and classified according to their Milo age-associated log-fold change as positive (logFC > 0.25) or negative (logFC < −0.25). Nuclei were then assigned to positive or negative comparison groups based on neighborhood membership, and nuclei represented in both groups were excluded. For each sample × group combination, log-normalized expression was aggregated by taking the mean across nuclei. Differential expression between positive and negative macrophage neighborhood groups was then performed using limma^90^ (v3.64.1), with donor-level replication accounted for using *duplicateCorrelation*, followed by linear modeling and empirical Bayes shrinkage under the fixed-effects model:

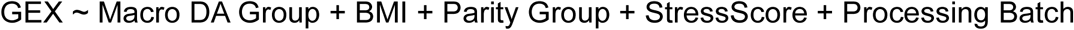

where GEX denotes aggregated log-normalized gene expression and Macro DA Group indicates membership in the positive (enriched with age) or negative (depleted with age) macrophage neighborhood group. Benjamini–Hochberg-adjusted P values were used for significance assessment, and moderated t-statistics for the macrophage neighborhood-group coefficient were used for downstream pathway analysis.

To assess whether macrophage neighborhood-level differential expression captured signals beyond those identified when macrophages were analyzed as a single annotated cell type, moderated t-statistics from the age-enriched versus age-depleted macrophage neighborhood comparison were compared with those from the corresponding whole-cell-type age analysis.

#### Pathway analyses

To address distinct analytic goals in different statistical settings, three pathway-analysis strategies were used: (1) downstream of per-cell-type DGE, (2) downstream of Milo neighborhood differential abundance analysis, and (3) descriptively to characterize distinct stromal states.

#### Pathway analyses for per-cell-type DGE

Pathway analysis was performed on the *dreamlet* differential expression fits using *zenith_gsa*, which extends the limma *camera* competitive gene set framework to *dreamlet* mixed-model results while accounting for inter-gene correlation. Hallmark gene sets from MSigDB^91,92^ (v2025.1.Hs) were used for analyses of Age and Parity, using the default inter-gene correlation of 0.01. Pathway significance was assessed using the BH adjusted FDR returned by *zenith_gsa* for each cell type–predictor analysis. For visualization, a pathway-level standardized effect statistic was calculated as the effect size divided by its standard error (delta / se); this value was displayed as the pathway Enrichment Score in figures and corresponds to the pathway-level t-statistic from *zenith_gsa*.

To identify genes driving pathway effects that were concordantly increased across cell types with positive age-associated enrichment, cell types with significant positive pathway enrichment (FDR ≤ 0.05, delta > 0) were first selected. Gene-level moderated t-statistics for genes within the pathway were then extracted from the corresponding *dreamlet* results. Consensus pathway drivers were defined as genes present in at least 75% of included cell types, positive in at least 80% of included cell types, and with mean gene-level moderated t-statistic ≥ 1 across included cell types. Up to 30 of the top genes meeting these criteria were visualized for the Age analysis, and the corresponding Parity gene-level t-statistics were displayed for the same genes and cell types.

#### Pathway analysis of Milo-defined macrophage neighborhoods

To characterize biological programs distinguishing age-enriched versus age-depleted macrophage neighborhoods, moderated t-statistics from the donor-blocked limma differential expression analysis were analyzed using limma *cameraPR*^93^ with Reactome gene sets from MSigDB (extracted by msigdbr v25.1.1). Gene sets were mapped to the analyzed gene universe and filtered to retain pathways containing 5–500 genes. *cameraPR* was run with the default inter-gene correlation of 0.01, and significance was assessed using the resulting Benjamini–Hochberg-adjusted FDR values. For figure display, a biologically informative subset of pathways was selected to summarize coherent inflammatory, regulatory, and stress-response programs.

#### Descriptive pathway scoring to characterize stromal states

To characterize transcriptionally distinct stromal states, Hallmark (MSigDB_Hallmark_2020) and Reactome (Reactome_Pathways_2024) gene sets retrieved from Enrichr, together with a recently reported 32-gene spatially derived p16-associated ovarian signature (BuckSenOvary) from postmenopausal human ovaries (Supplementary Note 1)^16^, were used for descriptive scoring. For Hallmark and Reactome analyses, genes with non-zero expression in at least 1% of nuclei in at least one stromal state were retained to define the expressed-gene universe. Each gene set was intersected with this universe, and only gene sets containing 25–300 retained genes were kept. Broad disease-, variant-, and infection-related Reactome wrapper pathways were excluded prior to scoring to improve interpretability. Per-nucleus pathway scores were then computed on log-normalized expression using PyUCell *compute_ucell_scores*, summarized as the median across nuclei for each donor– stromal state combination, and averaged across donors to yield donor-robust stromal-state pathway scores. For visualization, pathway specificity for each stromal state was defined relative to the mean score across stromal states. A curated subset of state-informative pathways was then displayed after clipping negative specificity values and applying per-pathway min–max scaling.

The BuckSenOvary signature was scored separately using the same UCell framework. Because two genes in the reported signature (*HLA-B* and *HLA-DRB1*) were not represented in the probe panel used for snucRNA-seq, only detected signature genes were retained for scoring. BuckSenOvary scores were summarized as donor-level medians for each cell type and displayed as boxplots across donors. Full pathway/signature selection, scoring, labeling, and plotting code is provided in the deposited repository.

#### Cell–cell communication analyses

To infer cell–cell communication programs, we used LIANA+^53^ (v1.5.0) followed by latent factor extraction with Tensor-cell2cell^54^ (v0.8.4). Cell types were first retained if they were represented by at least 10 nuclei in at least 10% of samples. Cell–cell communication was then inferred independently within each sample, grouping nuclei by annotated cell type and treating each tissue piece as a separate sample. LIANA+ was run using its default consensus ligand–receptor resource on the log-normalized expression matrix. A minimum of 5 nuclei was required for each cell type–sample combination, and an expression threshold of 10% was used internally by LIANA when scoring ligand–receptor interactions. Because all ligand–receptor pairs were retained for downstream analyses, interactions failing this threshold were not discarded but instead assigned the lowest ranks.

Per-sample LIANA results were then assembled into a four-dimensional tensor spanning sample × ligand–receptor pair × sender cell type × receiver cell type, using the LIANA consensus magnitude rank as the interaction score. Ligand–receptor pairs were retained in the tensor if they were present in at least 25% of samples, and missing values were represented as NaN. Tensor-cell2cell decomposition was then performed to derive 10 latent communication factors.

To interpret factor-specific communication programs, cell–cell connection strengths were examined using adjacency thresholds of 0.05, 0.08, and 0.1; these thresholds were used only to aid interpretation and visualization and do not represent statistical significance cutoffs. Ligand–receptor pairs distinguishing individual factors were identified by z-scoring factor loadings across factors, followed by manual curation of descriptive ligand–receptor groups for display to reduce redundancy.

To assess associations between sample-level latent communication factor scores and donor metadata, ordinary least-squares models with donor-clustered standard errors were fit using the design equation:

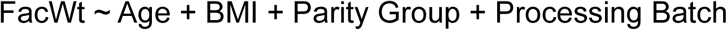

where FacWt denotes the sample-level latent communication factor score. Repeated samples from the same donor were accounted for using a cluster-robust sandwich estimator with clustering on Donor ID. StressScore was excluded for this analysis due to potential technical–biological confounding (see Supplementary Note 6). Benjamini– Hochberg correction was applied across factor results within each model term.

#### Statistics and reproducibility

Statistical analyses were performed as described in the corresponding sections. Repeated sampling from the same donor was accounted for using random effects, donor blocking, or donor-clustered standard errors where appropriate. Frequentist P values were obtained from the default tests or model-based statistics implemented in the indicated software packages unless otherwise stated. Bayesian compositional analyses with sccomp used the package’s default priors and inference settings unless otherwise specified, including default mean-prior settings for intercepts and coefficients and default overdispersion–mean association priors; primary analyses used Pathfinder inference and sensitivity analyses used Hamiltonian Monte Carlo inference. sccomp results were summarized using posterior effect estimates, credible intervals, posterior evidence, and Bayesian false-discovery rates rather than frequentist P values. Multiple-testing correction was applied within each analysis using the Benjamini–Hochberg procedure, graph-overlap spatial false-discovery correction, Bayesian false-discovery control, or the native false-discovery framework of the corresponding method, as appropriate. Variance inflation analyses indicated low collinearity among model covariates (adjusted GVIFs < 2), supporting their joint inclusion without evidence of problematic multicollinearity.

All computational analyses were run with fixed random seeds where applicable, and full code for preprocessing, modeling, visualization, and downstream analyses has been deposited with the study to support reproducibility.

#### Large language model usage

Under the direction of the authors, large language models (LLMs) were used for code generation and refining of written content to improve communication for the benefit of the reader. All content produced by LLMs was reviewed by the authors to ensure their intent was appropriately conveyed. The authors take full responsibility for all content submitted.

## Data availability

The raw and processed data reported in this study have been deposited in the Gene Expression Omnibus (GEO) repository under accession number GSE329417.

Bulk RNA-seq data from the GTEx v10 release is available from the open access GTEx portal (https://www.gtexportal.org/home/downloads/adult-gtex/bulk_tissue_expression) and the controlled access donor phenotypes and sample attributes are available under dbGaP accession number phs000424.v10.p2.

## Code availability

End-to-end code used for preprocessing, analysis, and visualization has been deposited to GitHub. The corresponding bioinformatic pipeline visualization is available in the repository and in Supplementary Figure 11. Additionally, an interactive data explorer application has been made available to support data reuse with a representative screenshot in Supplementary Figure 12.

GitHub repository: https://github.com/josef-byrne/snucRNA-seq_Ovary_Aging_Atlas

Hugging Face interactive data explorer app: https://huggingface.co/spaces/josef-byrne/postmenopausal-ovary-atlas-browser

## Acknowledgements

This was made possible by funding from the National Institutes of Health U54 AG075932 (PI: B.Sch.) and UH3 CA268105 (PI: F.E.D.) as part of the SenNet Consortium (https://sennetconsortium.org/) and a National Institute on Aging T32 AG052374 (USC Geroscience Training in the Biology of Aging) predoctoral fellowship to J.B.

## Author information

### Contributions

B.Sch. and F.E.D. conceived and supervised the study. E.S., M.E.G.P., and P.R.D. coordinated and performed clinical tissue collection and processing. B.So. contributed to receipt, handling, and storage of ovarian tissue samples. N.M. led, executed, and coordinated single-nucleus RNA-seq processing, with assistance from J.B. and T.T.. J.B. performed all computational analyses and figure generation and drafted the initial manuscript. M.A.W., B.Sch., and F.E.D. provided substantial intellectual input and detailed critical review that contributed to refinement of the manuscript. B.So. and N.M. contributed to data interpretation. B.Sch., F.E.D., and J.B. acquired funding. All authors reviewed and approved the manuscript.

