## Supplementary figures and images for "Single-nucleus profiling reveals age-associated remodeling opposed by parity in the postmenopausal human ovary"

### Extended Data Figure 1

#### Extended Data Fig. 1

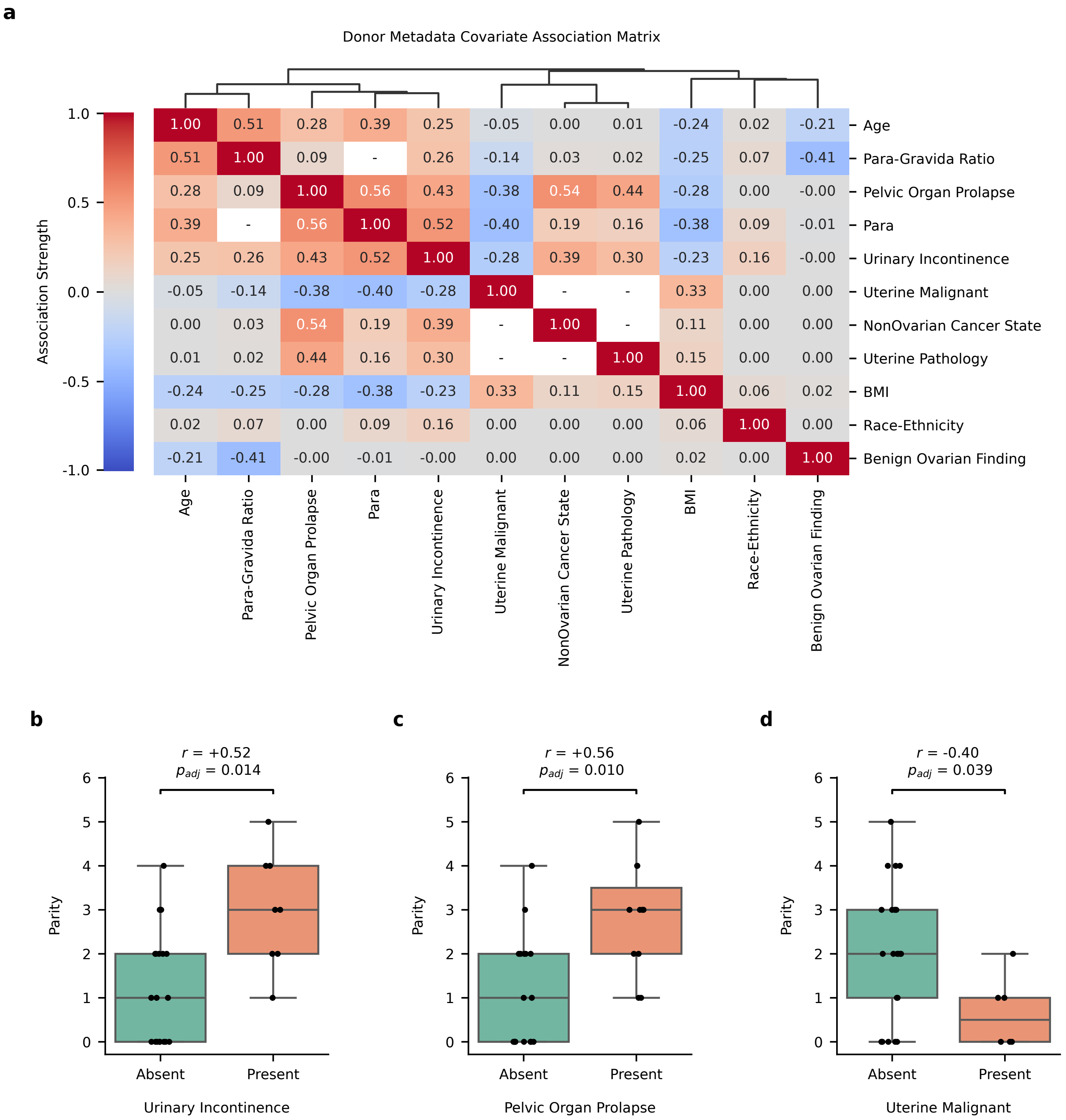

### Extended Data Figure 2

Extended Data Fig. 2

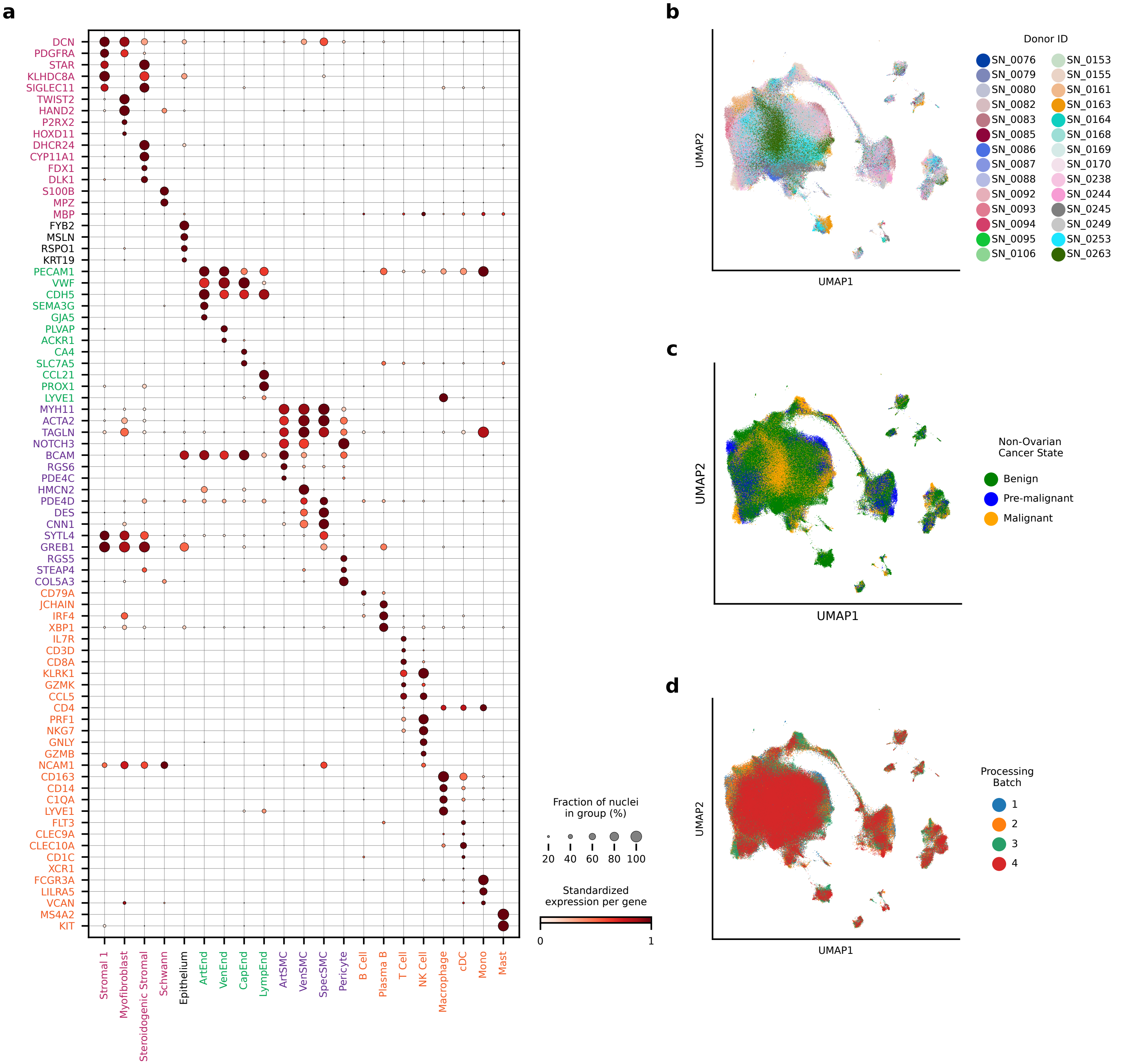

### Extended Data Figure 3

Extended Data Fig. 3

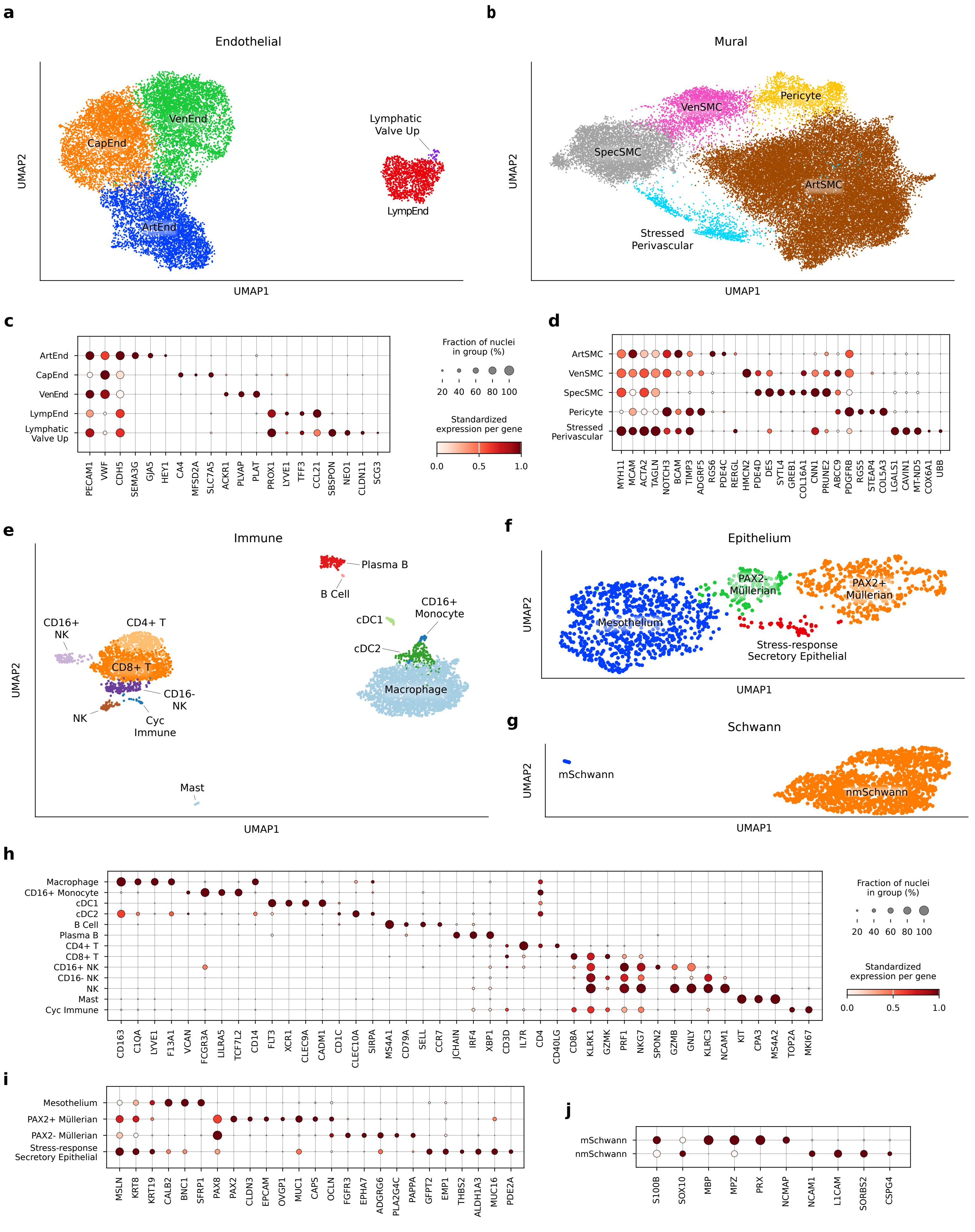

### Extended Data Figure 4

Extended Data Fig. 4

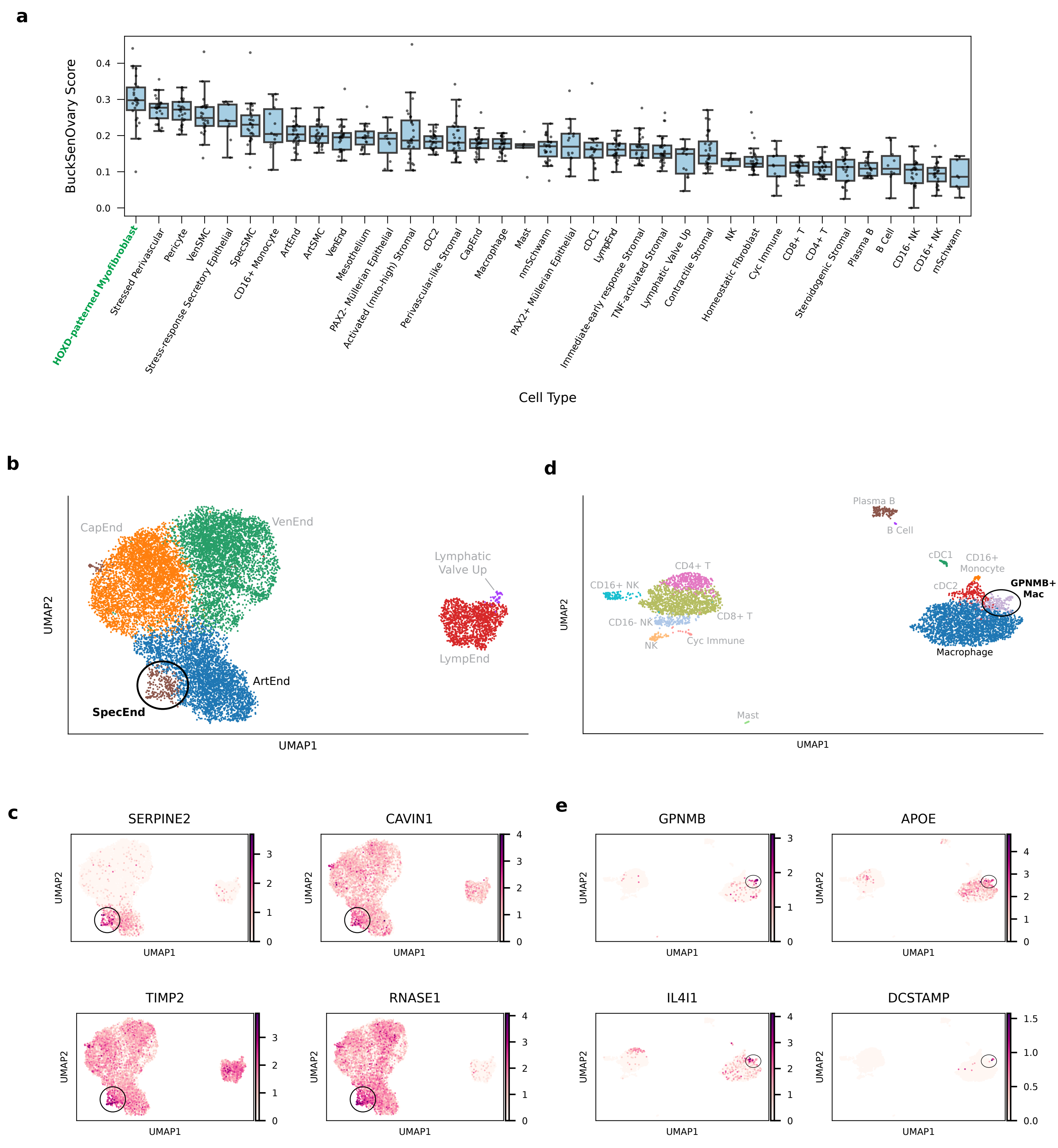

### Extended Data Figure 5

Extended Data Fig. 5

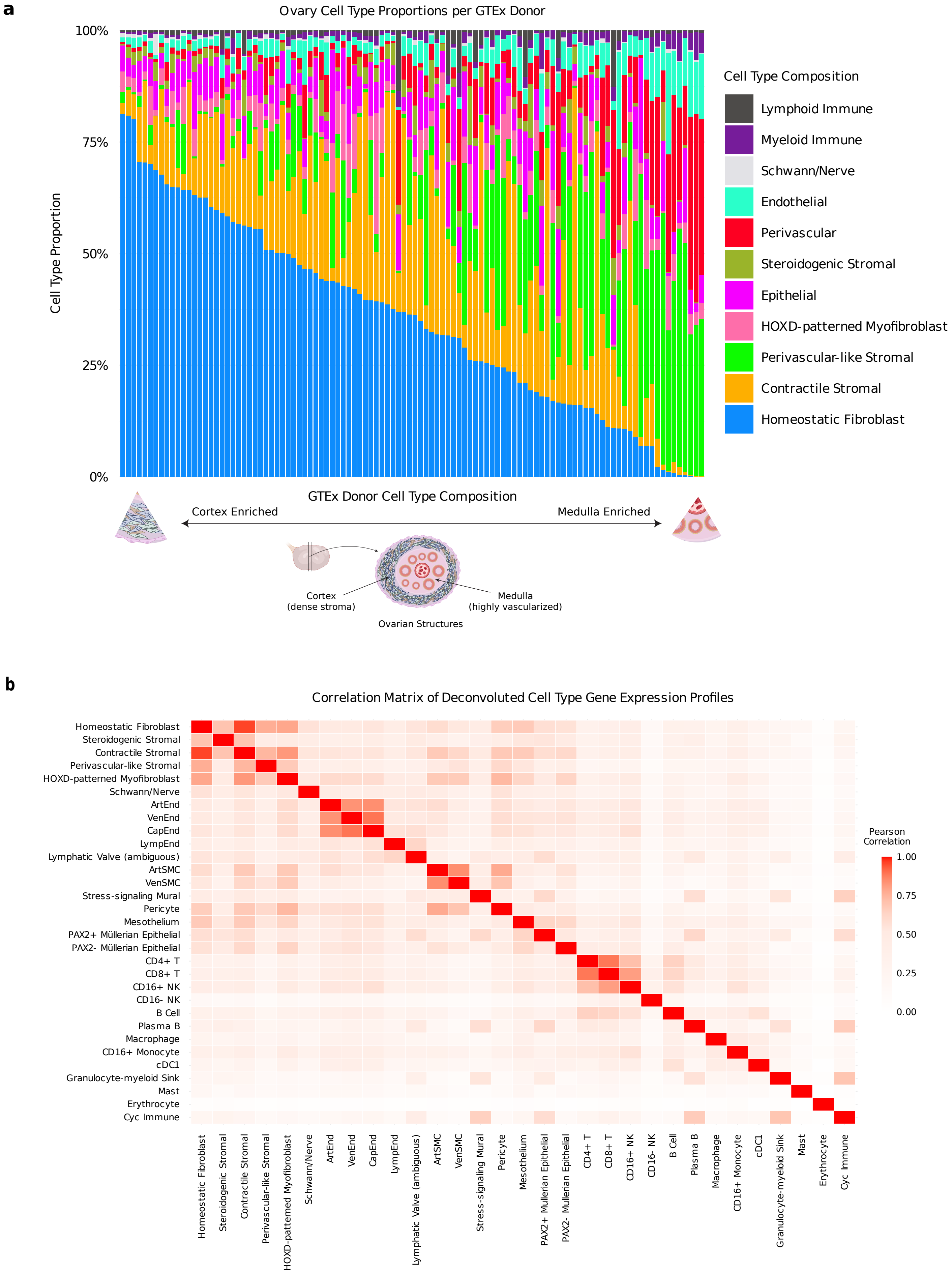

### Extended Data Figure 6

#### Extended Data Fig. 6

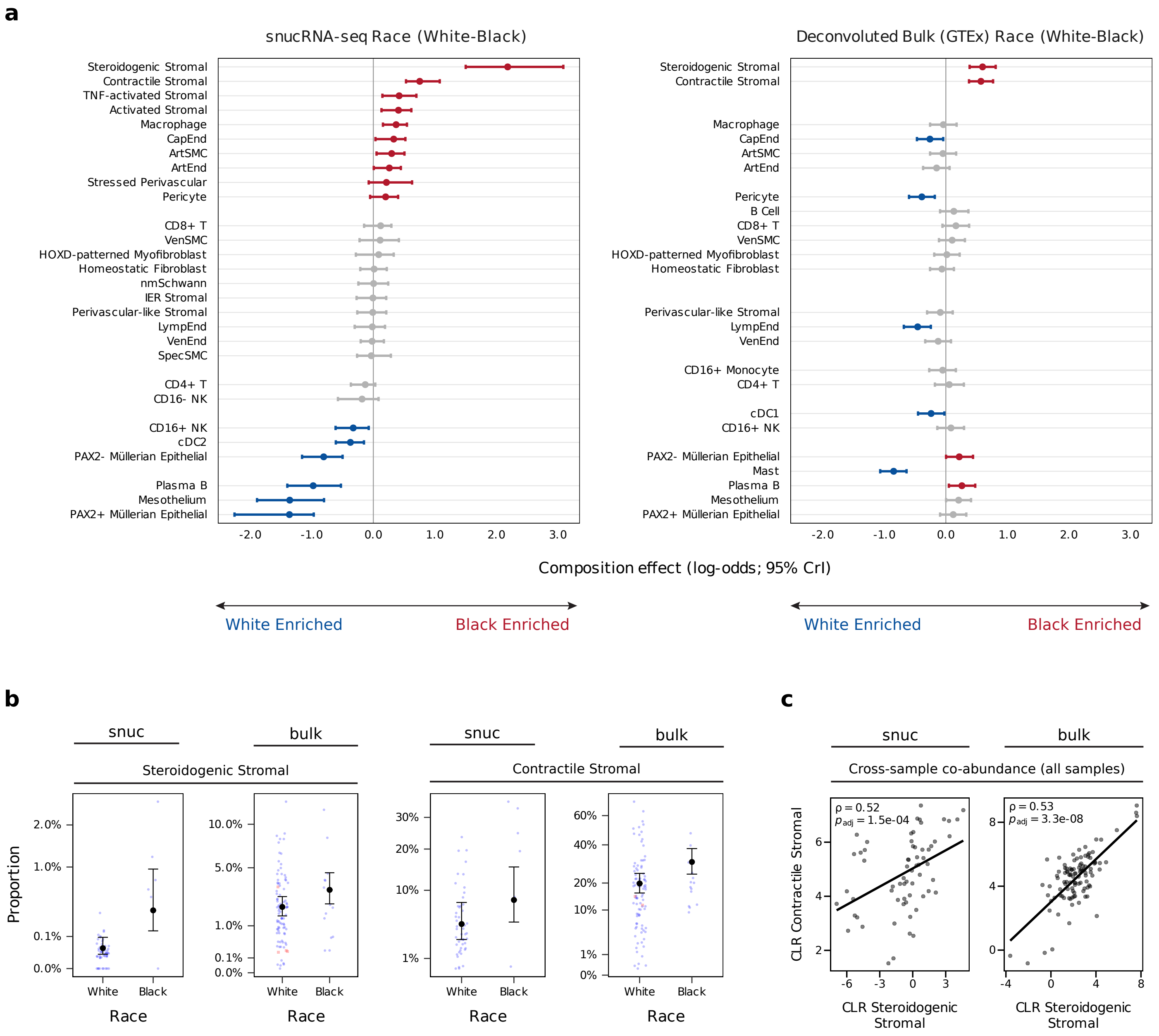

### Extended Data Figure 7

Extended Data Fig. 7

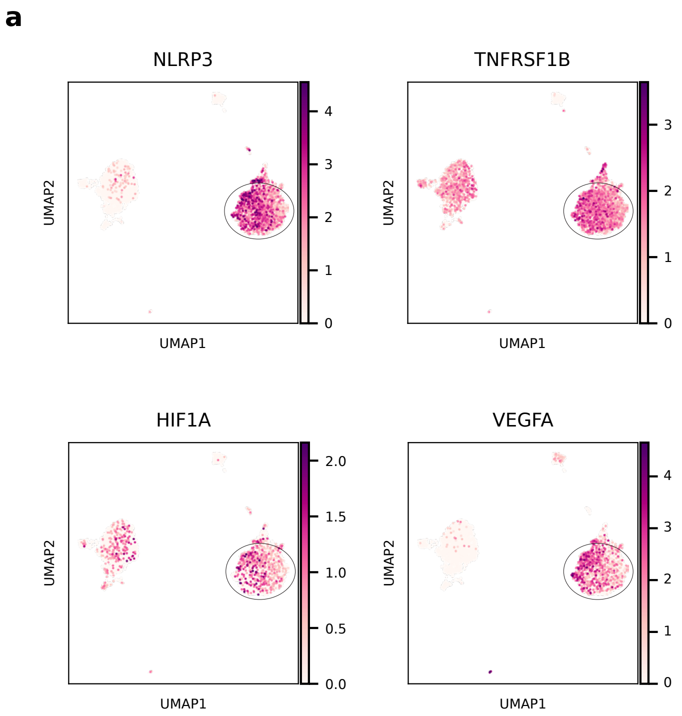


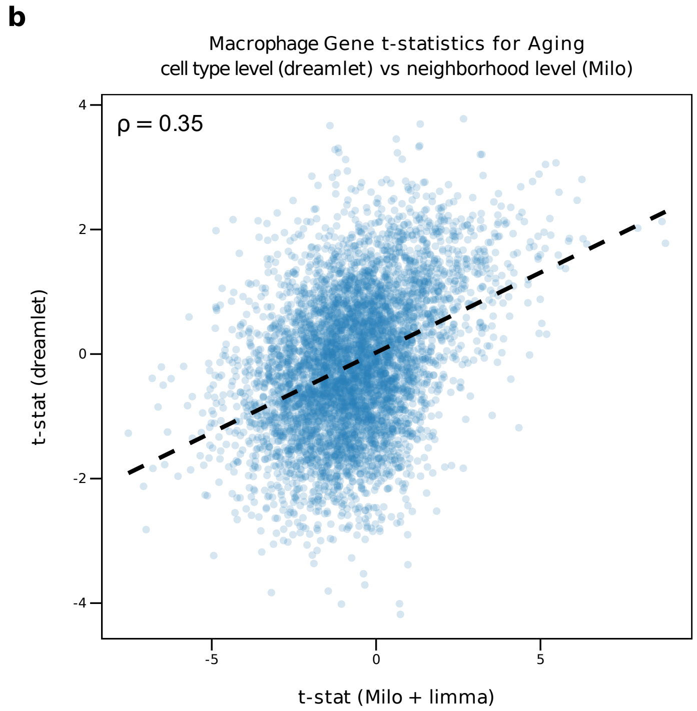

### Extended Data Figure 8

Extended Data Fig. 8

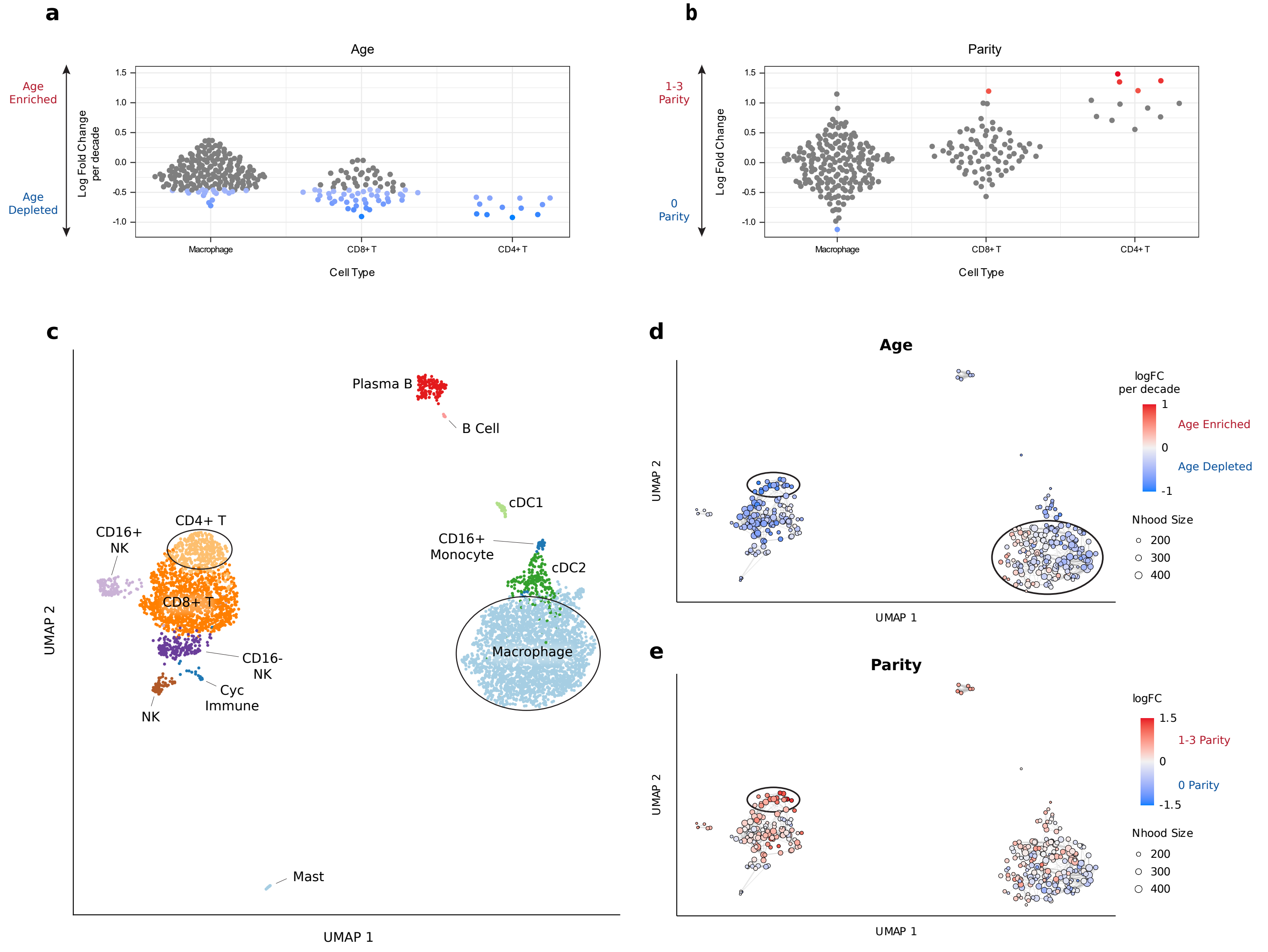

### Extended Data Figure 9

Extended Data Fig. 9

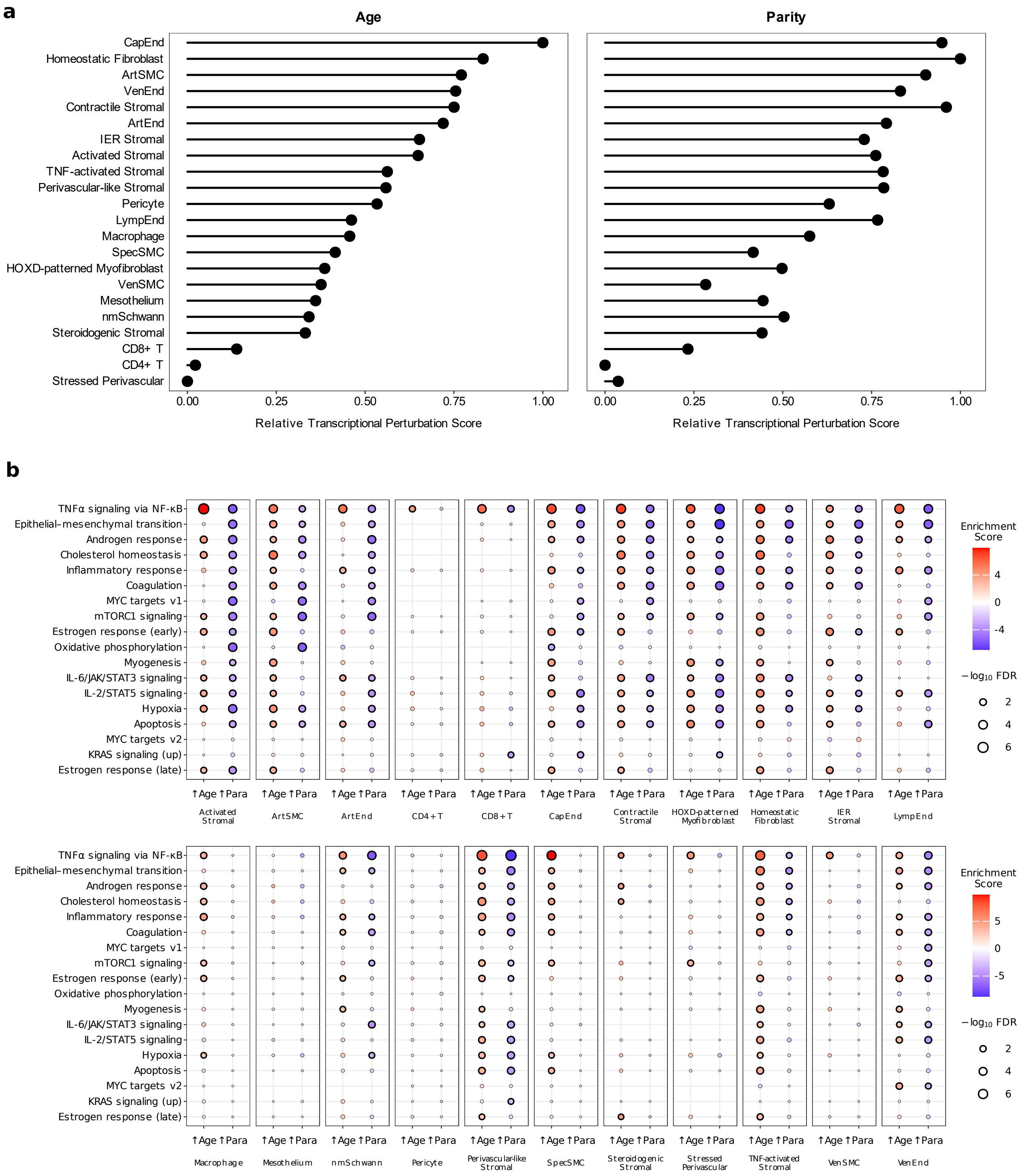

### Extended Data Figure 10

Extended Data Fig. 10

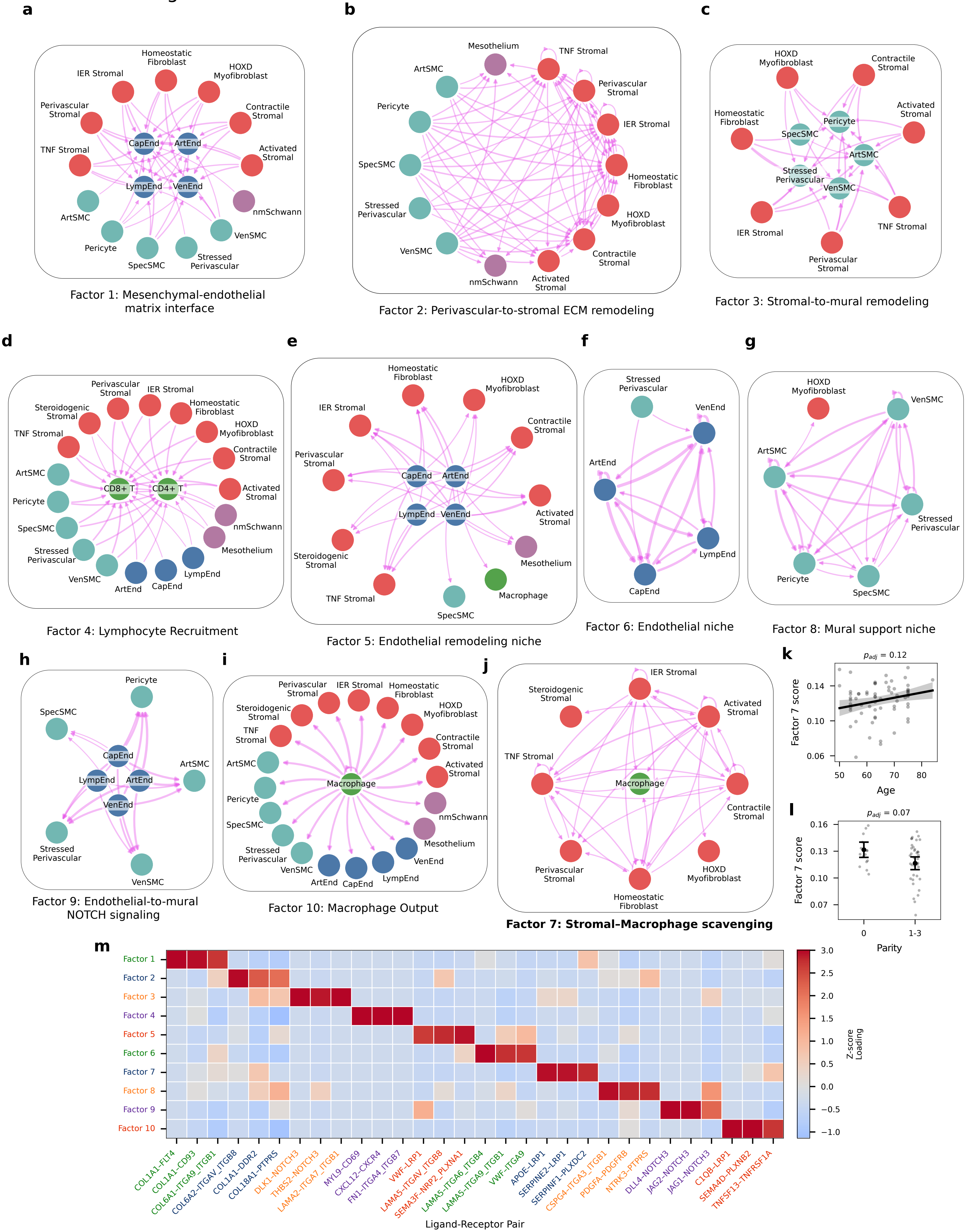
