## Supplementary Information for "Single-nucleus profiling reveals age-associated remodeling opposed by parity in the postmenopausal human ovary"

### Supplementary Notes

#### Supplementary Note 1. BuckSenOvary signature used for descriptive cell state characterization.

To compare atlas-defined cell states with a recently reported spatial p16-associated ovarian aging program, we used the 32-gene BuckSenOvary signature from Watson et al.[1]: *MYC*, *CCND2*, *TXNIP*, *RASD1*, *PKIG*, *EGR3*, *ATXN1*, *C3*, *CFD*, *HLA-B*, *HLA-DRB1*, *CD74*, *CD44*, *SRGN*, *COTL1*, *COL1A1*, *COL4A1*, *PCOLCE*, *SPARC*, *CCN1*, *CCN2*, *NID1*, *TAGLN*, *PDLIM7*, *CSRP1*, *TGFB1*, *IGFBP7*, *SAT1*, *CD9*, *PABIR1*, *KIAA1614* and *CTSB*. Because *HLA-B* and *HLA-DRB1* were not represented in the 10x Flex probe panel used for snucRNA-seq, these genes were excluded before scoring. The retained 30-gene signature was scored on log-normalized expression using PyUCell and summarized as described in Methods.

#### Supplementary Note 2. Immediate-early response signature used for sample-level StressScore.

To estimate transcriptional stress responses that arise during tissue collection prior to freezing and fixation, we used the “All Shared” immediate-early response gene set from Supplementary Table 21 of Marsh et al.[2] From this tab, we retained genes that were shared across at least two datasets comparing dissociation- or postmortem-associated stress responses across murine brain microglia[2], brain non-microglial macrophages[2,3], muscle stem cells[4], and kidney[5] cells. This resulted in the following stress-associated signature: *ATF3*, *DUSP1*, *EGR1*, *FOS*, *GEM*, *HSP90AA1*, *HSPA1A*, *HSPA1B*, *IER5*, *JUN*, *JUNB*, *JUND*, *KLF2*, *NFKBIZ*, *RHOB*, *ZFP36*. Although this signature was derived from the overlap of murine studies, strong concordance with postmortem-associated stress in human tissue has been reported[2].

**Supplementary Note 3. Review and curation of GTEx deconvolution annotations.**

GTEx ovary bulk RNA-seq samples were deconvolved using atlas-derived expression profiles. Because deconvolution profiles derived from fine single-nucleus states can be difficult to interpret in bulk tissue, we reviewed each deconvolved profile before reporting cell type results. Annotation review was based on top marker genes for each deconvolved expression profile, UCell scoring against canonical and atlas-derived marker signatures, and correlation patterns among deconvolved profiles.

This review supported most atlas-derived labels, but identified several profiles requiring relabeling, merging, or cautious interpretation. The profile originally labeled SpecSMC retained mural/smooth muscle marker support but was dominated by stress/chaperone-associated markers, including *DNAJA4*, *BAG3*, *HSPB1*, *CRYAB*, *HSPH1*, *SERPINH1*, *DNAJB1*, *HSPA1A* and *HSPA6*. Therefore, this profile was relabeled Stress-signaling Mural rather than interpreted as a distinct specialized smooth muscle cell population. The profile originally labeled cDC2 showed a mixed myeloid/granulocyte-like marker pattern, including *PADI4*, *S100A12*, *CXCR2*, *CSF3R*, *FCGR3B*, *S100A8*, *MMP9* and *TREM1*, and was relabeled Granulocyte-myeloid Sink to avoid over interpreting it as a conventional dendritic cell population. The profile originally labeled Lymphatic Valve Up showed partial endothelial/lymphatic marker-score support, but its top markers included a mixed set of developmental, neural/neuroendocrine and epithelial-associated genes, including *ZIC2*, *ZIC5*, *IRX1*, *IRX2*, *SCG3*, *GRIK1*, *NTS*, *CACNA1E*, *GRIN2B*, *CEACAM6*, *S100P* and *PKP1*. Because this profile was not supported by a clean lymphatic valve marker pattern, we relabeled it Lymphatic Valve (ambiguous). Finally, the mSchwann and nmSchwann profiles both showed Schwann/nerve-associated signal, but their separation was not considered robust in GTEx bulk deconvolution. The mSchwann and nmSchwann profiles were distinguishable by their top marker genes but were merged for reporting as Schwann/Nerve because both represented nerve/Schwann-associated signals in GTEx bulk RNA-seq. The mSchwann profile was dominated by myelin-associated Schwann markers, including *MBP*, *NCMAP*, *FA2H*, *CLDN19*, *PMP2*, *UGT8*, *MPZ* and *S100B*, while also containing neural/nerve-associated transcripts such as *SHISA7*, *CAMK2B*, *LGI1*, *KIF1A*, *SV2B*, *SLC17A7* and *HCN1*. The nmSchwann profile included Schwann-associated genes such as *SOX10*, *CDH19*, *S100B*, *ERBB3* and *GFRA3*, together with broader peripheral nerve/neural-associated transcripts including *NRXN1*, *GRIK2*, *GRIK3*, *SLC5A7*, *CNTN2*, *CACNG5*, *LRRTM1*, *NTM* and *CADM2*. Because bulk deconvolution cannot reliably distinguish discrete ovarian Schwann states from broader peripheral nerve transcript content, these profiles were merged and not interpreted separately.

Relabeled and merged profiles were retained in the deconvolution output and sccomp compositional testing to preserve the full modeled composition and avoid redistributing ambiguous signal across other profiles. However, transcriptionally ambiguous or nuisance profiles (e.g. erythrocytes) were not emphasized in downstream biological interpretation or in the main reported GTEx validation results. Full top-marker tables, atlas marker scores, and deconvolved proportion summaries are provided in Supplementary Table 3.

### Supplementary Note 4. Descriptive parity-group patterns in raw cell-type proportions

Parity was modeled as a categorical variable in bins of nulliparous (0 births), moderate parity (1–3 births) and higher parity (4–5 births) because raw sample-level proportions suggested that parity-associated compositional differences were not uniformly linear across cell types. Several vascular and mural populations, including arterial endothelial, arterial smooth muscle, venous endothelial and venous smooth muscle cells, appeared highest in donors with 1–3 live births, whereas CD4+ and CD8+ T cell proportions showed a more graded increase across parity groups (**Supplementary Fig. 4**). Because these raw summaries are unadjusted for covariates and affected by compositional dependence, they were treated as descriptive support for parity-group modeling rather than as standalone evidence of nonlinear dose-response. Statistical inference was based on the multivariable sccomp model.

### Supplementary Note 5. Sensitivity of sccomp compositional results to posterior inference method.

Primary sccomp compositional analyses used Pathfinder inference, the default scalable inference method in sccomp. To evaluate sensitivity to posterior inference method, we refit the snucRNA-seq age and parity models using Hamiltonian Monte Carlo inference and summarized the resulting posterior distributions with half-eye plots (**Supplementary Figs. 8 and 9**). Hamiltonian Monte Carlo produced wider posterior uncertainty intervals and fewer Bayesian FDR significant effects, but posterior median effect directions were broadly concordant with Pathfinder for the major age- and parity-associated cell-type changes highlighted in the manuscript. These results support the qualitative robustness of the principal age- and parity-associated compositional findings, while indicating that formal significance calls were more conservative under Hamiltonian Monte Carlo inference.

### Supplementary Note 6. Assessment of sample-level StressScore effects on cell–cell communication results.

Sample-level StressScore was associated with several inferred communication factors; therefore, we interpreted stress-associated factors cautiously. However, Factor 7, which was highlighted in the main manuscript, showed no detectable association with StressScore, and its age-associated increase and parity-associated decrease were preserved after StressScore adjustment (Supplementary Table 7). These findings support interpretation of Factor 7 as a stromal–macrophage remodeling/scavenging communication program that is not primarily explained by sample-level tissue collection stress variation.

### Supplementary Figures

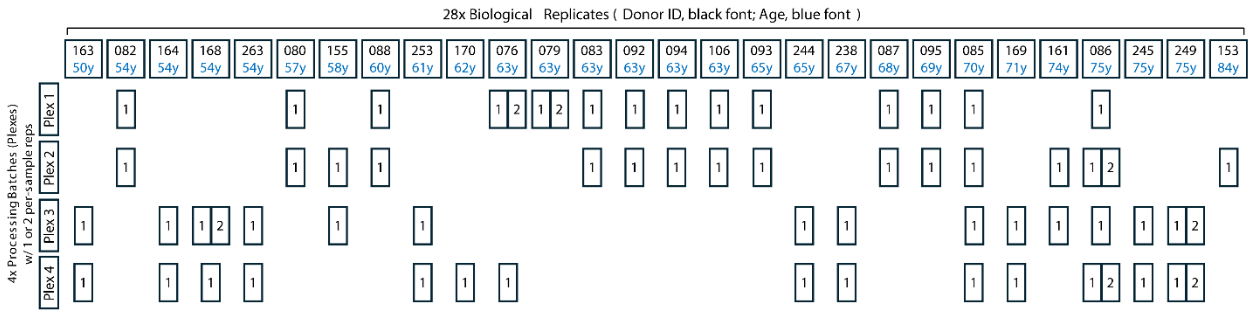


**Supplementary Fig. 1 | Experimental design for multiplexed ovary snucRNA-seq profiling.** Schematic overview of donor age distribution and sample allocation across four 10x Flex processing plexes. Columns represent de-identified donors ordered by age; donor ID (SenNet ID) is shown in black and age in blue. Numbers within each plex row indicate the number of tissue samples from that donor processed in the corresponding plex.

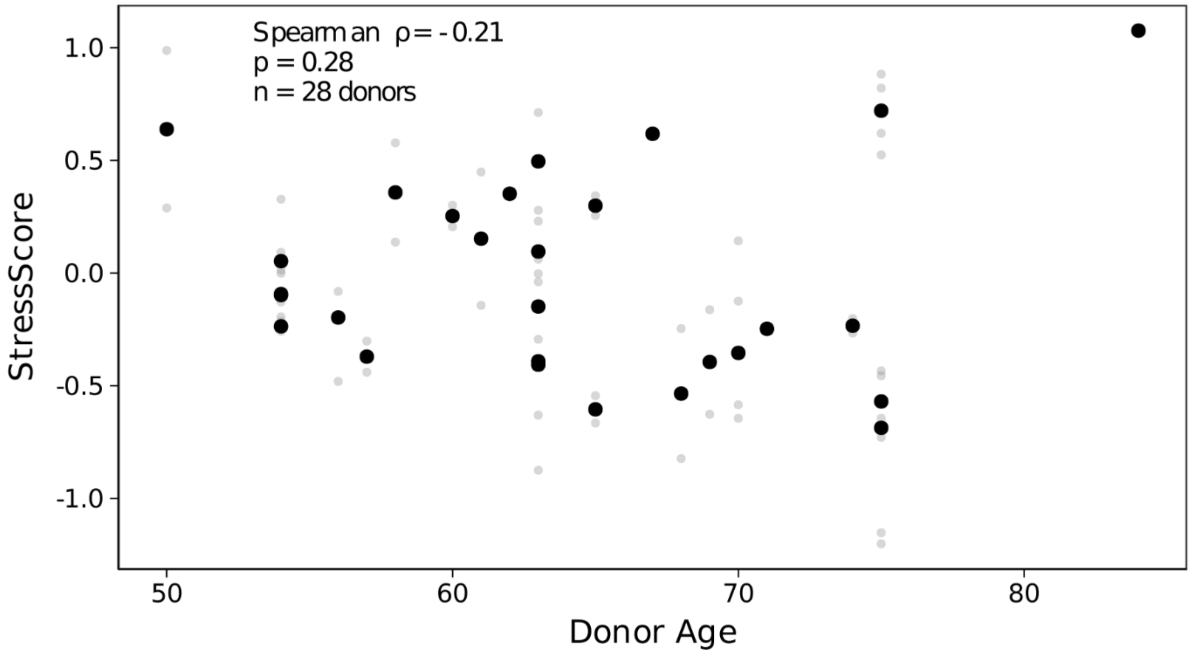


**Supplementary Fig. 2 | Relationship between donor age and StressScore.** Grey points show sample-level StressScore values used in downstream models; black points show donor-level medians used for correlation with donor age. Spearman correlation was calculated using donor-level median StressScore to avoid pseudoreplication from multiple samples per donor.

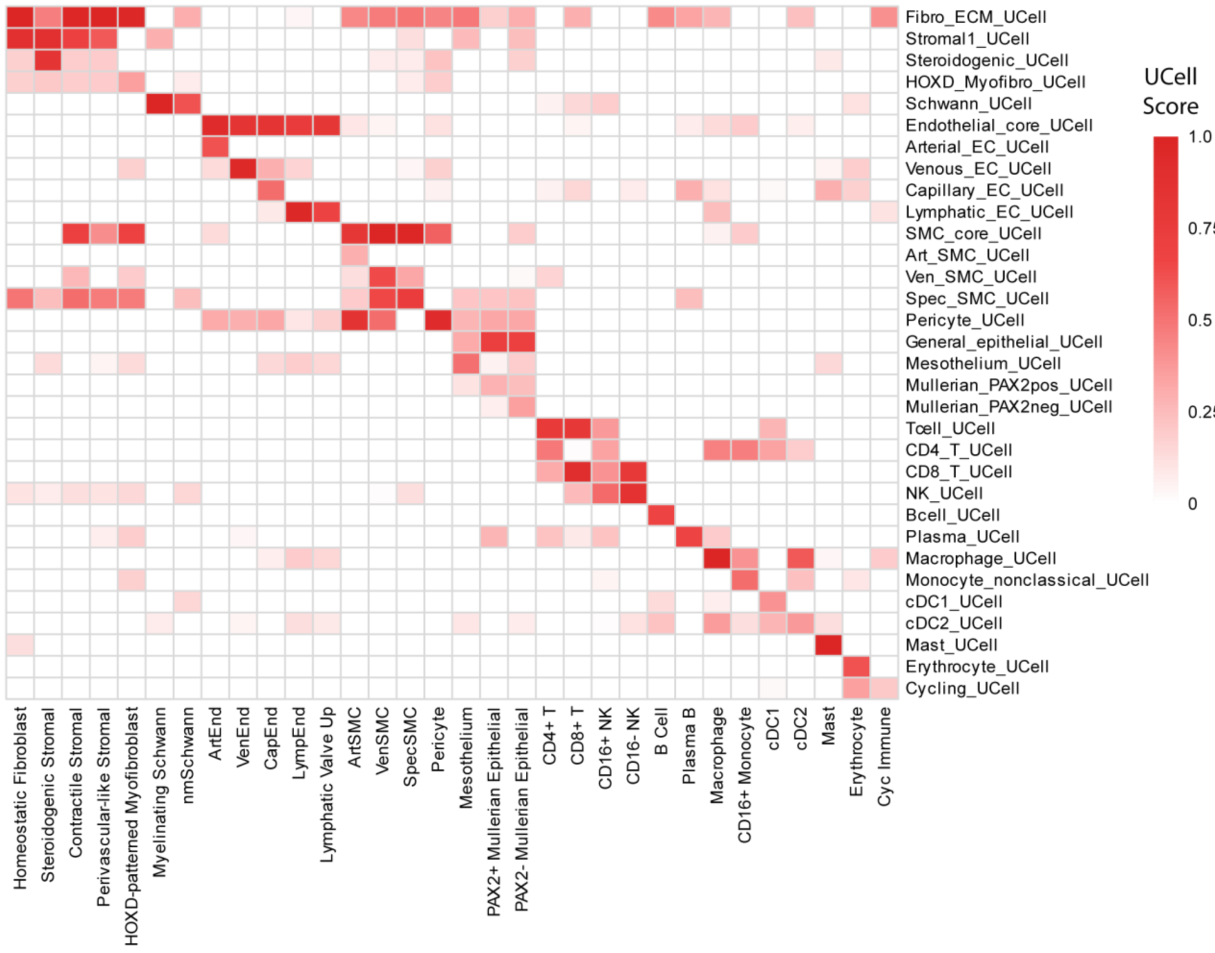


**Supplementary Fig. 3 | Atlas marker signature overlap across deconvolved cell-type expression profiles.** Heatmap showing UCell scores for curated atlas marker signatures across inferred deconvolved cell-type mean expression profiles. Higher scores indicate stronger overlap between a deconvolved profile and the corresponding atlas-derived marker signature.

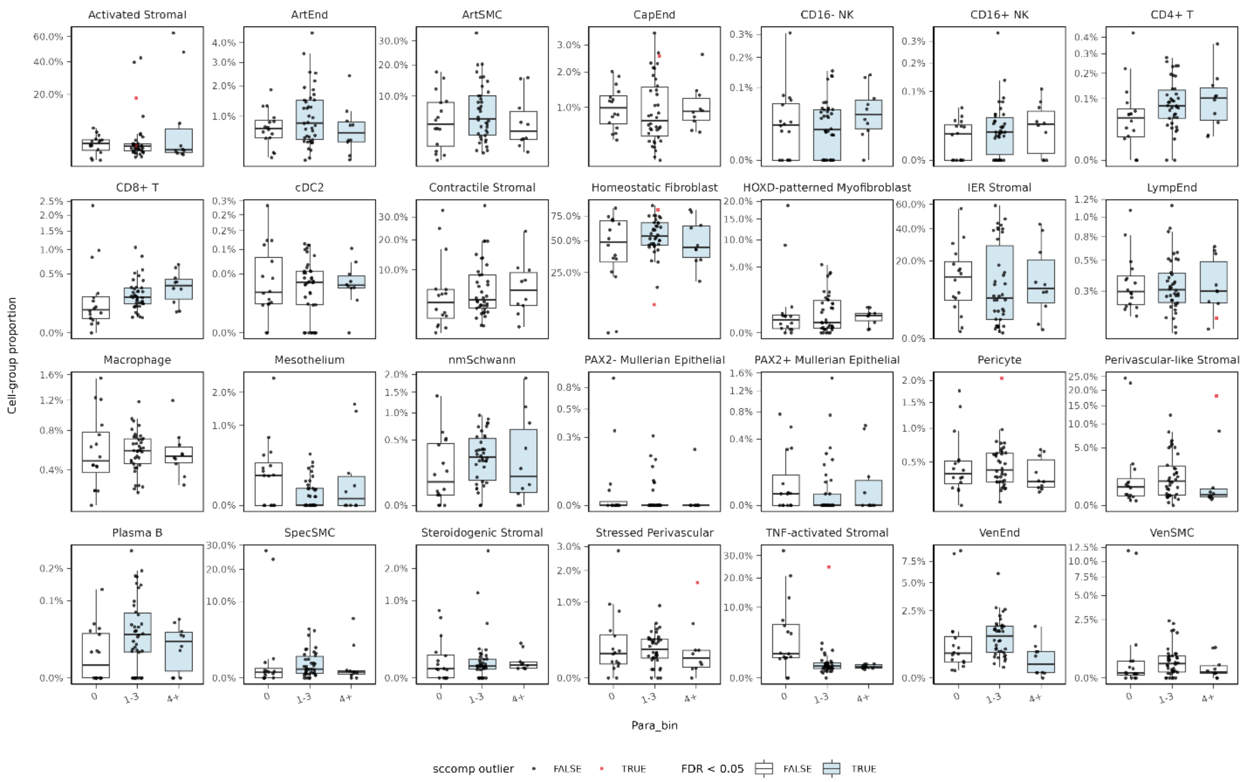


**Supplementary Fig. 4 | Raw snucRNA-seq cell type proportions by parity group.** Boxplots show observed sample-level proportions for each fine cell type tested in the sccomp compositional analysis. Black dots indicate observations retained by sccomp and red squares indicate sample–cell-type observations censored as outliers during model fitting. Blue boxes denote cell types with significant parity-associated composition effects in the multivariable sccomp model (Bayesian FDR ≤ 0.05; absolute composition effect >0.1 logit units). The fitted trends are from the multivariable sccomp model described in Methods; therefore, raw univariate trends may not directly mirror covariate-adjusted model estimates.

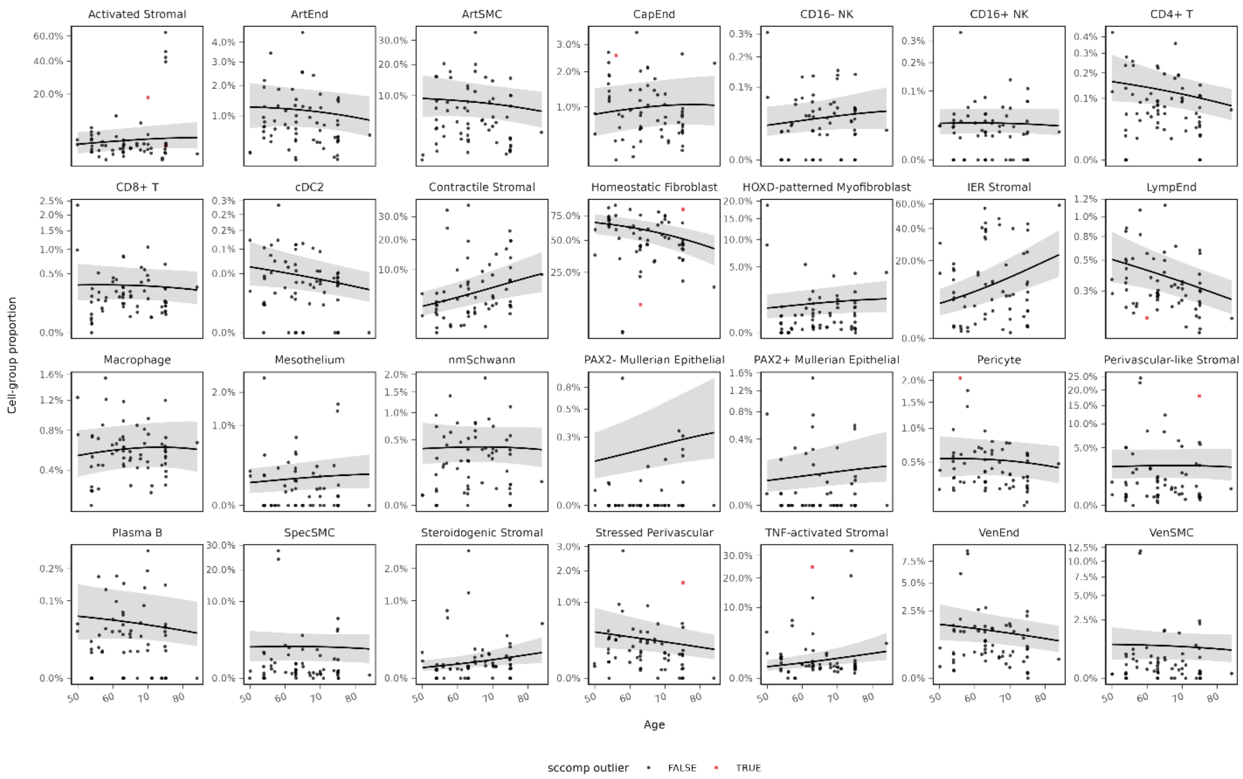


**Supplementary Fig. 5 | Raw snucRNA-seq cell-type proportions across age with sccomp marginal model fit.** Points show observed sample-level proportions for each fine cell type tested in the sccomp compositional analysis. Red squares indicate sample–cell-type observations identified as outliers and censored during sccomp model fitting. Black lines and grey ribbons show marginalized sccomp model-predicted proportions and 95% credible intervals across donor age. The multivariable model included age, parity group, BMI, processing plex, simplified race/ethnicity, sample-level StressScore and a donor-level random intercept; therefore, raw univariate trends may not directly mirror covariate-adjusted model estimates.

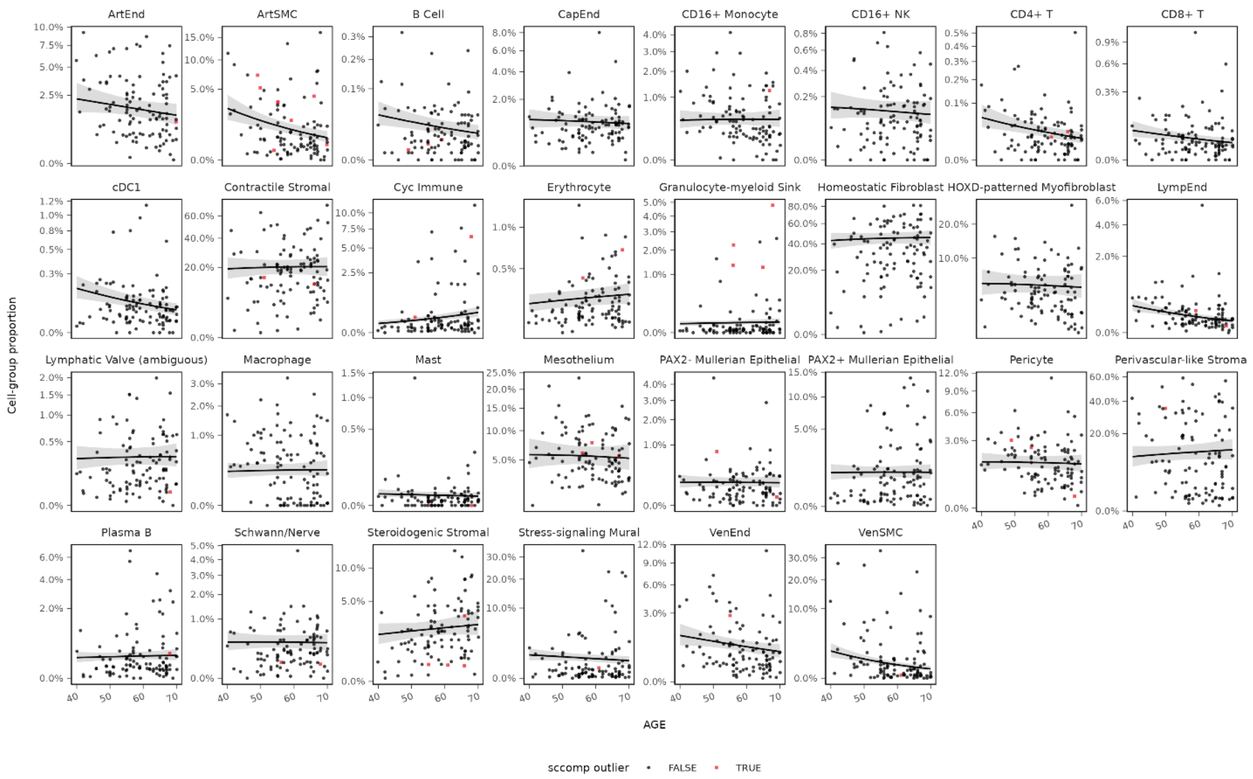


**Supplementary Fig. 6 | Raw GTEx deconvolved ovary cell-type proportions across age with sccomp marginal model fit.** Points show deconvolved sample-level proportions for each cell type tested in the GTEx sccomp compositional analysis. Red squares indicate sample–cell-type observations identified as outliers and censored during sccomp model fitting. Black lines and grey ribbons show marginalized sccomp model-predicted proportions and 95% credible intervals across donor age. The fitted trends are from the multivariable GTEx sccomp model described in Methods; therefore, raw univariate trends may not directly mirror covariate-adjusted model estimates.

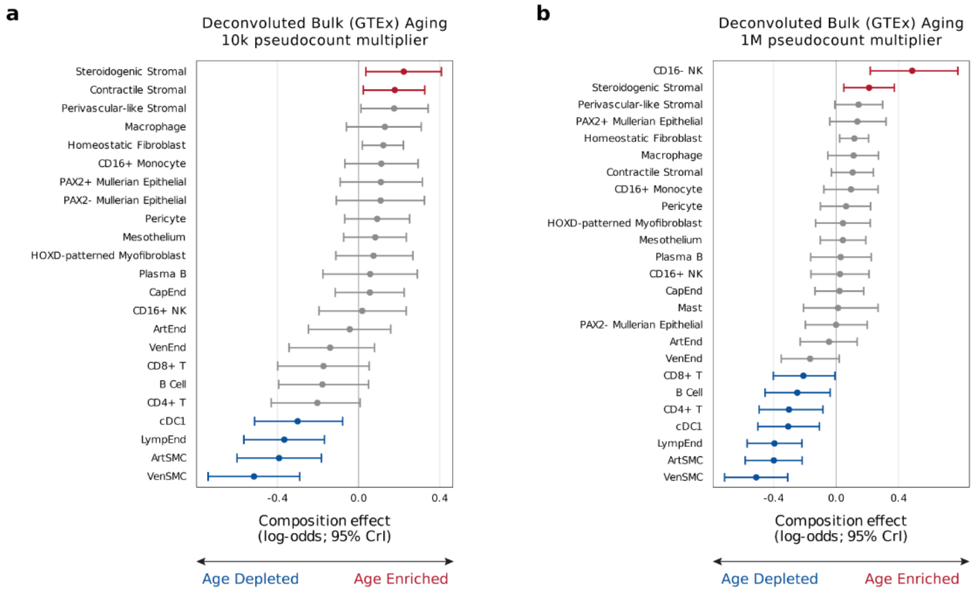


**Supplementary Fig. 7 | Sensitivity of GTEx deconvolved bulk sccomp age effects to pseudocount multiplier.**

Forest plots compare age-associated sccomp composition effects in GTEx deconvolved ovary bulk RNA-seq using alternative pseudocount multipliers of 10k (**a**) and 1,000,000 (**b**) for converting deconvolved proportions to integer counts. Points indicate posterior median composition effects and horizontal bars indicate 95% credible intervals. Positive values indicate age-enriched deconvolved cell-type proportions, and negative values indicate age-depleted proportions. Age-effect directions and major findings were broadly concordant across multiplier choices, with sensitivity concentrated among low-abundance cell types more affected by rounding to zero at lower pseudocount multipliers. The 100,000× multiplier was retained for the primary analysis.

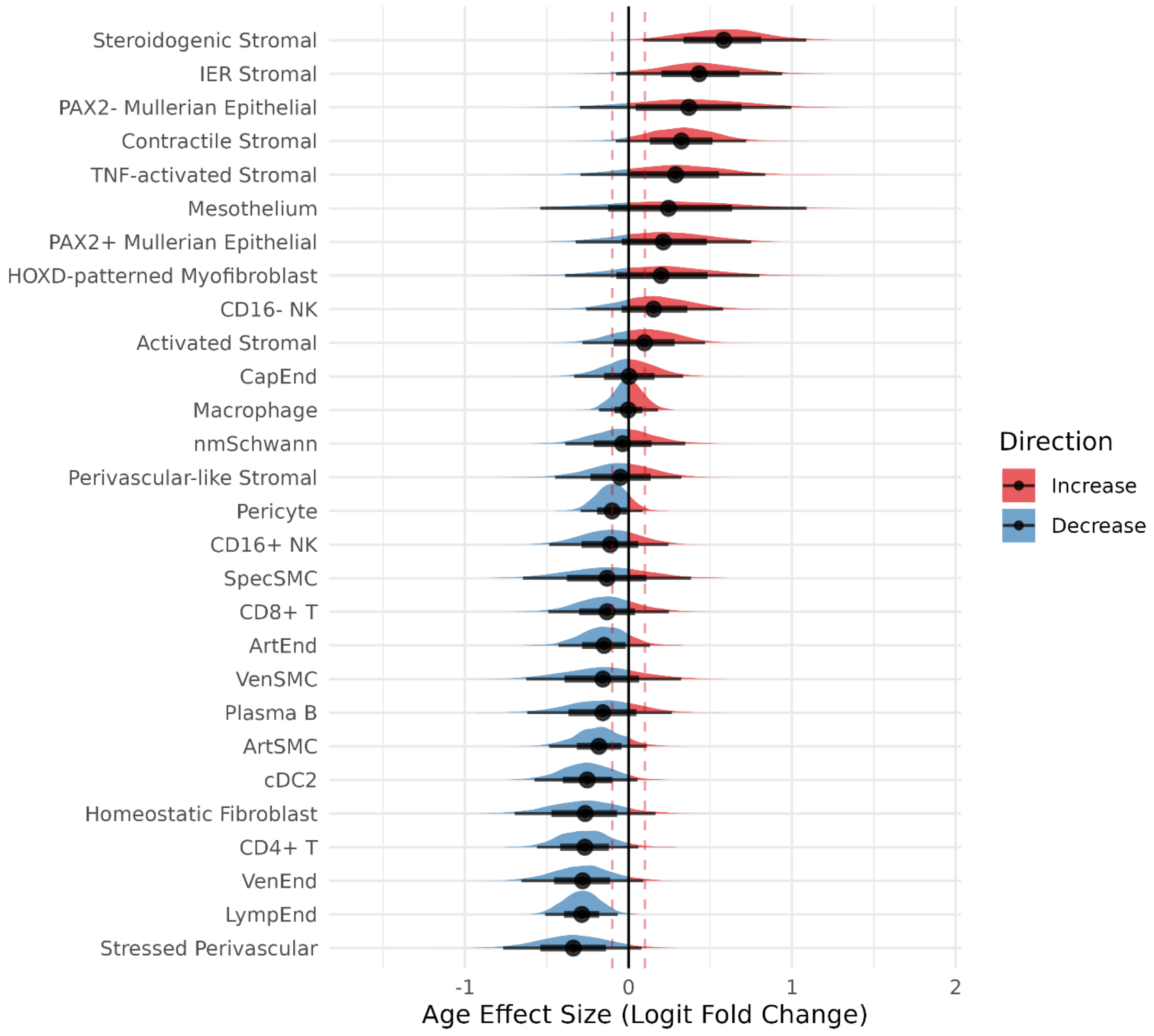


**Supplementary Fig. 8 | Hamiltonian Monte Carlo sensitivity analysis for snucRNA-seq compositional changes with age.**
Half-eye plots show posterior distributions of sccomp composition effect estimates for age across fine cell types from the snucRNA-seq dataset using Hamiltonian Monte Carlo inference. Slabs indicate posterior density, and intervals denote 66% and 95% credible intervals. Positive values indicate increasing relative abundance and negative values indicate decreasing relative abundance with age. The direction of the major effects was broadly concordant with the primary Pathfinder-based analysis, although posterior intervals were wider.

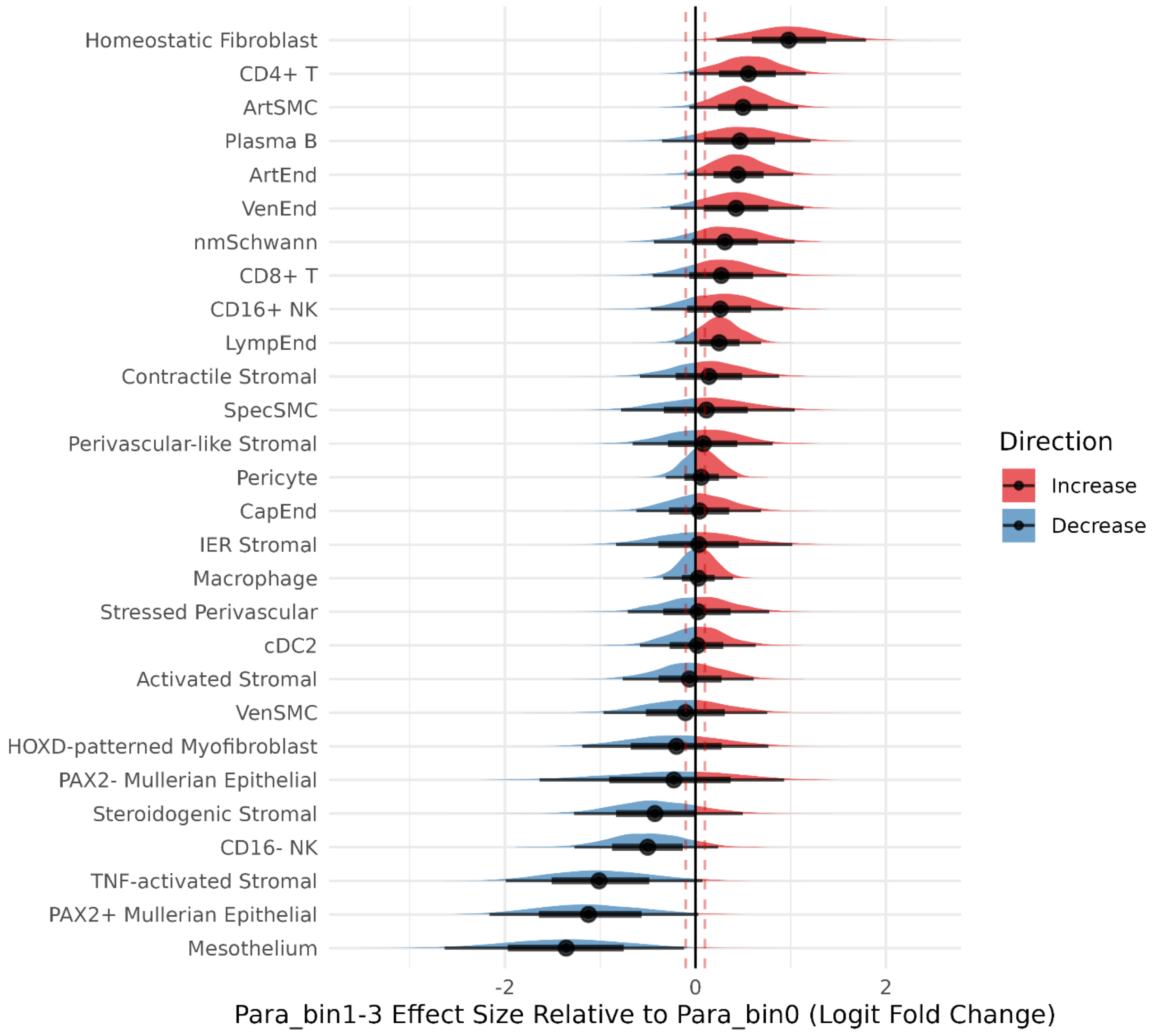


**Supplementary Fig. 9 | Hamiltonian Monte Carlo sensitivity analysis for snucRNA-seq compositional changes by parity group.**

Half-eye plots show posterior distributions of sccomp composition effect estimates for the parity contrast across fine cell types from the snucRNA-seq dataset using Hamiltonian Monte Carlo inference. Slabs indicate posterior density, and intervals denote 66% and 95% credible intervals. Positive values indicate relatively higher abundance in the contrasted parity group and negative values indicate relatively lower abundance. The direction of the major effects was broadly concordant with the primary Pathfinder-based analysis, although posterior intervals were wider.

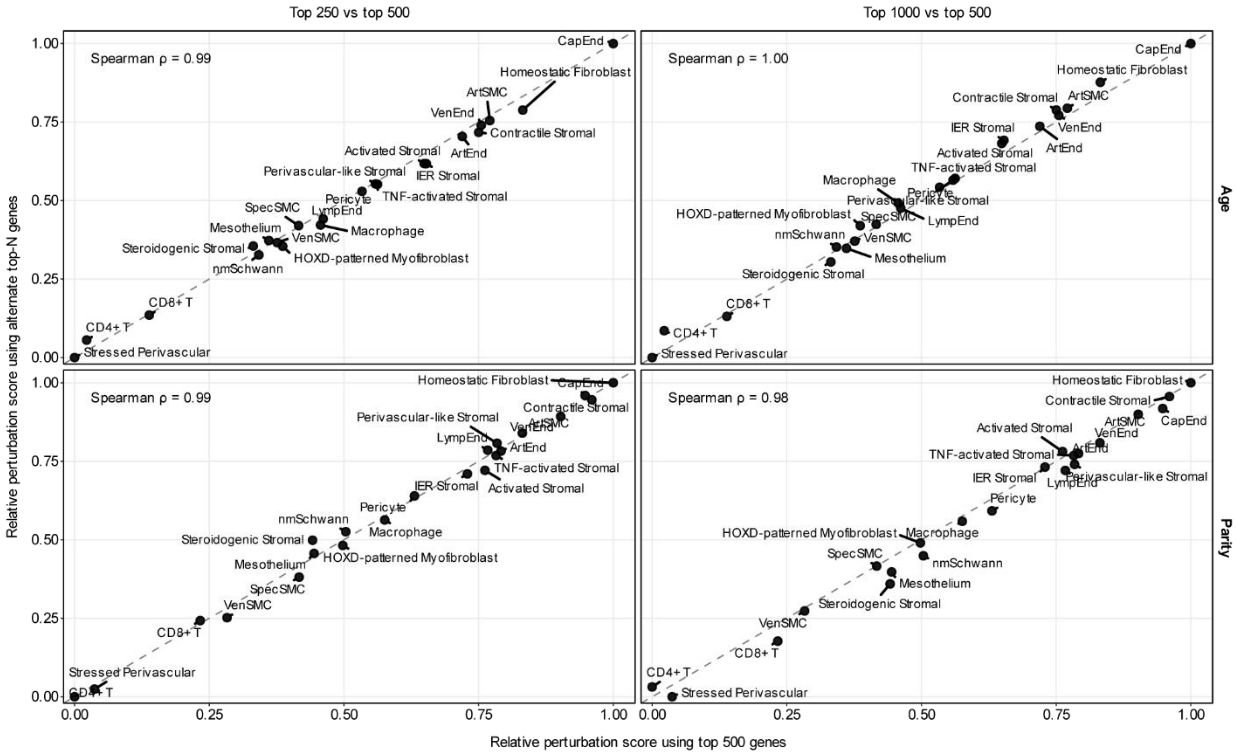


**Supplementary Fig. 10 | Sensitivity of transcriptional perturbation scores to the number of genes retained.** Relative perturbation scores were recomputed using the top 250, 500 or 1,000 genes ranked by absolute moderated t-statistic for each cell type and contrast. Scatterplots compare scores computed using 250 or 1,000 genes against the primary 500-gene score used in the main analysis. Dashed lines indicate identity; Spearman correlations are shown for each comparison. Parity denotes the 1–3 versus 0 live-birth contrast.

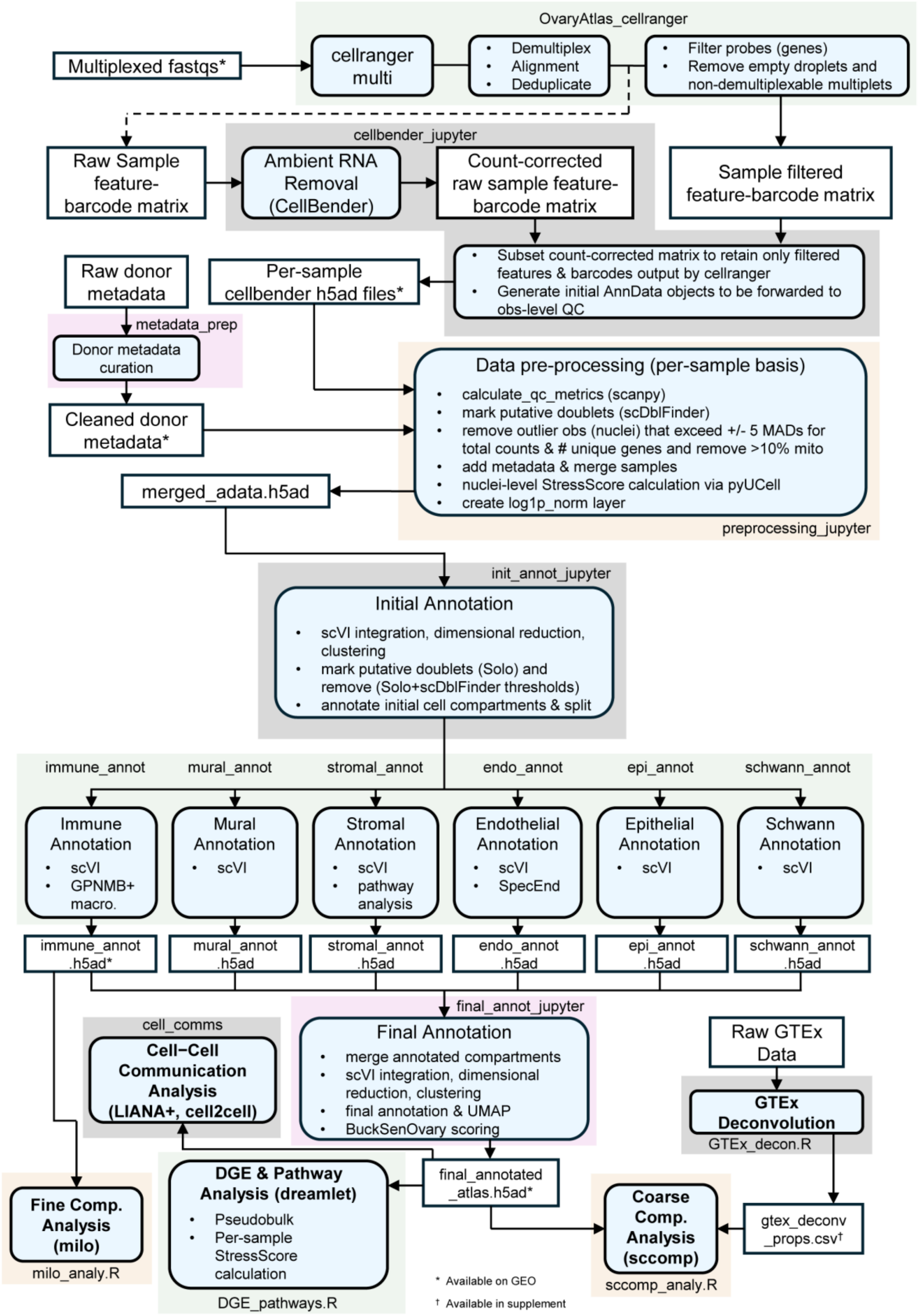
OvaryAtlas_cellranger
Multiplexed fastqs*
cellranger multi
Demultiplex
Alignment
Deduplicate
Filter probes (genes)
Remove empty droplets and non-demultiplexable multiplets
cellbender_jupyter
Raw Sample feature-barcode matrix
Ambient RNA Removal (CellBender)
Count-corrected raw sample feature-barcode matrix
Sample filtered feature-barcode matrix
Subset count-corrected matrix to retain only filtered features & barcodes output by cellranger
Generate initial AnnData objects to be forwarded to obs-level QC
Raw donor metadata
Per-sample cellbender h5ad files*
metadata_prep
Donor metadata curation
Cleaned donor metadata*
Data pre-processing (per-sample basis)
calculate_qc_metrics (scanpy)
mark putative doublets (scDblFinder)
remove outlier obs (nuclei) that exceed +/- 5 MADs for total counts & # unique genes and remove >10% mito
add metadata & merge samples
nuclei-level StressScore calculation via pyUCell
create log1p_norm layer
preprocessing_jupyter
merged_adata.h5ad
init_annot_jupyter
Initial Annotation
scVI integration, dimensional reduction, clustering
mark putative doublets (Solo) and remove (Solo+scDblFinder thresholds)
annotate initial cell compartments & split
immune_annot
mural_annot
stromal_annot
endo_annot
epi_annot
schwann_annot
Immune Annotation
scVI
GPNMB+ macro.
Mural Annotation
scVI
Stromal Annotation
scVI
pathway analysis
Endothelial Annotation
scVI
SpecEnd
Epithelial Annotation
scVI
Schwann Annotation
scVI
immune_annot .h5ad*
mural_annot .h5ad
stromal_annot .h5ad
endo_annot .h5ad
epi_annot .h5ad
schwann_annot .h5ad
final_annot_jupyter
cell_comms
Cell–Cell Communication Analysis (LIANA+, cell2cell)
Final Annotation
merge annotated compartments
scVI integration, dimensional reduction, clustering
final annotation & UMAP
BuckSenOvary scoring
Raw GTEx Data
GTEx Deconvolution
GTEx_decon.R
Fine Comp. Analysis (milo)
milo_analy.R
DGE & Pathway Analysis (dreamlet)
Pseudobulk
Per-sample StressScore calculation
DGE_pathways.R
final_annotated _atlas.h5ad*
Coarse Comp. Analysis (sccomp)
sccomp_analy.R
gtex_deconv _props.csv†
* Available on GEO
† Available in supplement

**Supplementary Fig. 11 | Bioinformatic workflow for ovary snucRNA-seq atlas generation and analysis.** Overview of computational processing, quality control, annotation, and downstream analysis steps used to generate the final annotated atlas from multiplexed snucRNA-seq data. Major processing and analysis steps are shown as rounded light-blue boxes, scripts/notebooks are indicated by unboxed labels within colored workflow boundaries, and generated files are shown as white boxes. Asterisks denote files available through GEO; the dagger denotes output provided in the supplementary tables.

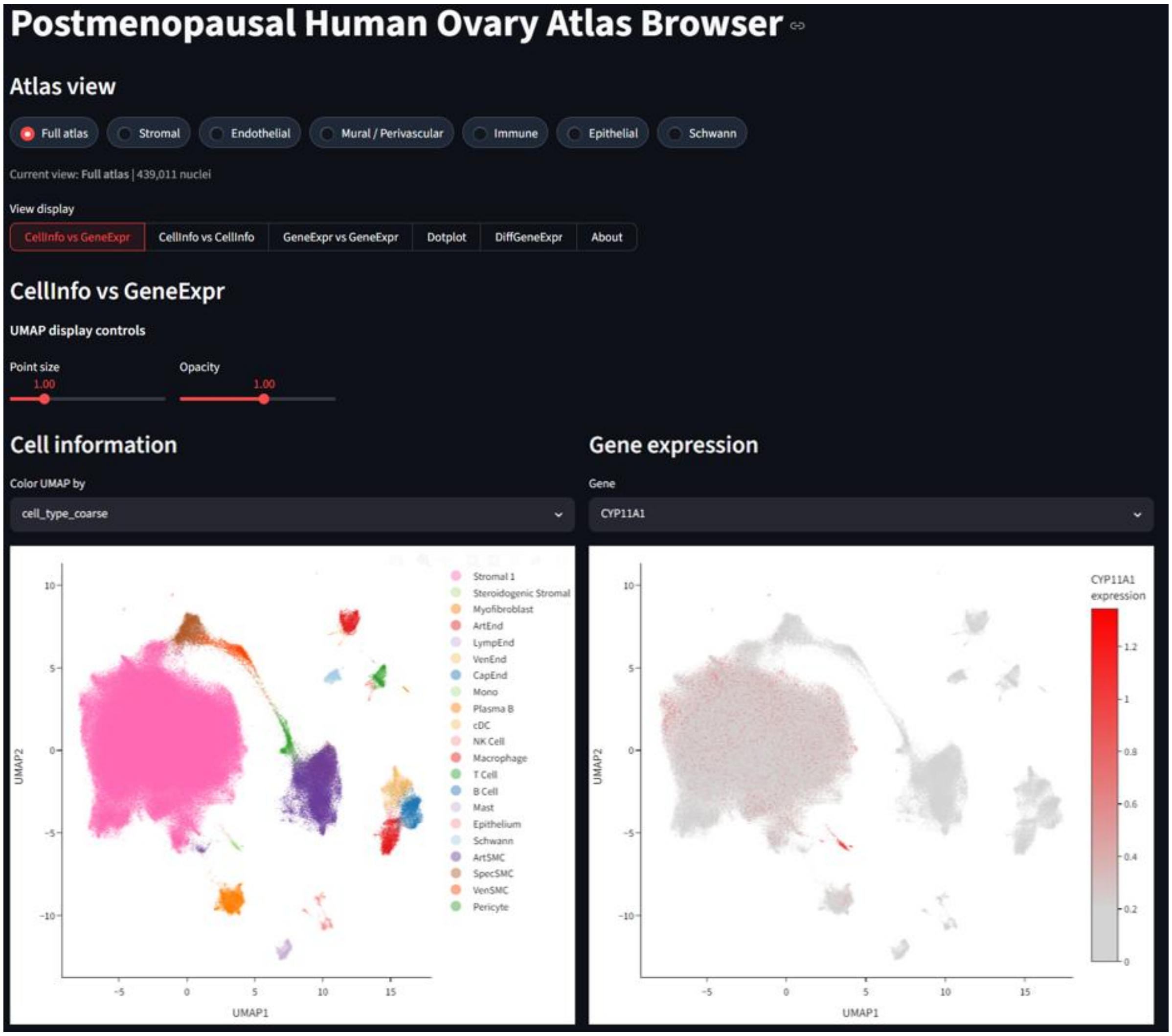


**Supplementary Fig. 12 | Interactive browser for the postmenopausal human ovary atlas.** Representative screenshot of the atlas browser, enabling exploration of cell annotations and gene expression across the full atlas or selected cellular compartments. Additional tabs provide comparisons of cell metadata and gene expression, dot plots, and differential gene expression results.[6]

### Supplementary Tables

**Supplementary Table 1. Study metadata and preprocessing quality control.**
(1) De-identified donor-level metadata, including age, BMI, parity, race/ethnicity, surgical indication, and clinical/pathology covariates. (2) Sample-level wet-lab processing information for the ovarian tissue samples profiled by snucRNA-seq. (3) Per-sample Cell Ranger/library quality-control metrics and downstream nucleus-level QC summaries, including scDblFinder doublet calls, 5-MAD count/gene outlier filtering, and mitochondrial transcript percentages. (4) Sample-level StressScore covariate used for downstream modeling, computed as the median robust z-scored immediate-early response score across cell types within each sample.

**Supplementary Table 2. Cell state marker genes and characterization.**
(1) Top 100 marker genes for each annotated fine cell state from whole-atlas comparisons against all other nuclei. (2) Top 100 marker genes for each annotated fine cell state from compartment-specific comparisons against other states within the same broad compartment. Marker genes were identified using scanpy *rank_genes_groups* with the Wilcoxon rank-sum test on log-normalized expression and filtered for Benjamini–Hochberg-adjusted $P \leq 0.05$, expression in ≥10% of nuclei in the target state, and positive log-fold change. (3) Stromal pathway scoring results used to characterize transcriptionally distinct stromal states. UCell pathway scores were summarized across donor–cell-state combinations and reported with state-specificity scores, display indicators, and relative enrichment values used for the Fig. 2d heatmap.

**Supplementary Table 3. GTEx ovary bulk RNA-seq deconvolution results.** (1) Top 50 marker genes inferred from deconvolved cell-type expression profiles, ranked by log2 fold-change relative to the average of all other deconvolved profiles. (2) Atlas marker gene signatures used to evaluate deconvolved cell-type identity. (3) UCell atlas marker-signature scores across deconvolved cell-type expression profiles. (4) Deconvolved cell-type proportions for GTEx ovary RNA-seq samples. (5) Summary statistics for deconvolved cell-type proportions across GTEx ovary RNA-seq samples, reported as percentages.

**Supplementary Table 4. Cell type compositional analysis results from snucRNA-seq and GTEx deconvolved bulk RNA-seq.** (1) Cell-type inclusion criteria for snucRNA-seq sccomp analysis. (2) Cell-type inclusion criteria for GTEx deconvolved bulk sccomp analysis. (3) Full sccomp model results for snucRNA-seq. (4) Full sccomp model results for GTEx deconvolved bulk RNA-seq. (5) Marginalized sccomp age predictions and fold changes relative to age 55. (6) Marginalized sccomp categorical predictions for parity and race/ethnicity contrasts. (7) Sample–cell-type observations censored as outliers during snucRNA-seq sccomp model fitting. (8) Sample–cell-type observations censored as outliers during GTEx deconvolved bulk sccomp model fitting. (9) Pairwise Spearman correlations between CLR-transformed cell-type proportions in snucRNA-seq and GTEx deconvolved bulk data.

**Supplementary Table 5. Milo immune neighborhood differential abundance and macrophage neighborhood characterization.** (1) Neighborhood-level quality control and inclusion metrics for the immune Milo analysis. Neighborhoods were retained for testing if they contained nuclei from ≥10 donors and ≥15 samples and if the three largest donor contributions accounted for ≤80% of neighborhood nuclei; all generated immune neighborhoods passed these filters. (2) Neighborhood-level differential abundance results for Age and Parity; Age was modeled per decade and Parity represents the 1–3 versus 0 live birth contrast. SpatialFDR reports graph-overlap-weighted FDR correction from miloR. (3) Cell type-level summaries of immune neighborhood differential abundance results. (4) Gene-level limma differential expression results comparing age-enriched and age-depleted macrophage neighborhoods. (5) Reactome cameraPR pathway results for age-enriched versus age-depleted macrophage neighborhoods. (6) Gene-level statistics for selected macrophage pathways highlighted in the manuscript.

**Supplementary Table 6. Differential gene expression and pathway analysis results.** (1) Gene-level dreamlet differential expression results for the primary Age and Parity contrasts across annotated cell types. P values were adjusted within each cell type/assay using Benjamini–Hochberg correction. (2) Cell type-level transcriptional perturbation scores and sensitivity analysis, computed as the root mean square of moderated t-statistics among the top 250, 500 or 1,000 genes ranked by absolute t-statistic; the primary analysis used the top 500 genes. (3) Hallmark pathway results from zenith/camera for the primary Age and Parity contrasts across annotated cell types.

**Supplementary Table 7. Cell–cell communication analysis results.** (1) Cell-type inclusion metrics for LIANA+/Tensor-cell2cell analysis, including total nuclei, sample representation, donor contribution, and inclusion status for primary cell–cell communication analysis. (2) Sample-level cell2cell context factor scores for the ten inferred communication factors. (3) Regression results associating sample-level communication factor scores with donor/sample covariates using ordinary least squares models with donor-clustered standard errors. (4) Sensitivity regression analysis results including sample-level StressScore as a covariate. (5) Top 50 ligand-receptor pairs associated with each communication factor, with raw cell2cell loadings shown across all factors. (6) Z-scored ligand-receptor pair loadings across factors for highlighted ligand-receptor pairs used to interpret communication factor identities. (7) Cell-type sender and receiver connectivity counts across communication factors and edge-weight thresholds. (8) Thresholded sender-receiver edge list for factor-specific communication networks.

6. Byrne, J. Postmenopausal Human Ovary Atlas Browser. https://huggingface.co/spaces/josef-byrne/postmenopausal-ovary-atlas-browser.
